# Consumption of human-relevant levels of sucrose-water rewires macronutrient uptake and utilization mechanisms in a tissue specific manner

**DOI:** 10.1101/2024.08.31.610015

**Authors:** Saptarnab Ganguly, Tandrika Chattopadhyay, Rubina Kazi, Souparno Das, Bhavisha Malik, M.L. Uthpala, Padmapriya S. Iyer, Mohit Kashiv, Anshit Singh, Amita Ghadge, Shyam D. Nair, Mahendra S. Sonawane, Ullas Kolthur-Seetharam

**Affiliations:** Tata Institute of Fundamental Research, Hyderabad, Telangana 500046, India; Centre for innovation in molecular and pharmaceutical sciences, Dr. Reddy’s Institute of Life Sciences, Hyderabad, Telangana 500046, India; Department of Biological Sciences, Tata Institute of Fundamental Research, Mumbai, Maharashtra 400005, India; Advanced Research Unit on Metabolism, Development and Aging (ARUMDA), TIFR, India; Centre for DNA Fingerprinting and Diagnostics, Hyderabad, Telangana 500039, India

**Author notes:** Address for correspondence : Prof. Ullas Kolthur-Seetharam - Tata Institute of Fundamental Research, Hyderabad, Telangana 500046, India. Contact number, email id, Prof. Mahendra S. Sonawane - Department of Biological Sciences, Tata Institute of Fundamental Research, Mumbai, Maharashtra 400005, India. Contact number, email id. SG, TC : Equal contribution. SD, BM : Equal contribution.

**Keywords:** Insulin resistance, gut transporters, systems physiology, mitochondria, nutrient sensing, gene expression

## Abstract

Consumption of sugar-sweetened beverages (SSBs) have been linked to metabolic dysfunction, obesity, diabetes and enhanced risk of cardiovascular diseases across all age-groups globally. Decades of work that have provided insights into pathophysiological manifestations of sucrose overfeeding have employed paradigms that rarely mimic human consumption of SSBs. Thus, our understanding of multi-organ cross-talk and molecular and/or cellular mechanisms, which operate across scales and drive physiological derangement is still poor. By employing a paradigm of sucrose water feeding in mice that closely resembles chronic SSB consumption in humans (10% sucrose in water), we have unraveled hitherto unknown tissue-specific mechanistic underpinnings, which contribute towards perturbed physiology. Our findings illustrate that systemic impaired glucose homeostasis, mediated by hepatic gluconeogenesis and insulin resistance, does not involve altered gene expression programs in the liver. We have discovered the pivotal role of the small intestine, which in conjunction with liver and muscles, drives dyshomeostasis. Importantly, we have uncovered rewiring of molecular mechanisms in the proximal intestine that is either causal or consequential to systemic ill-effects of chronic sucrose water consumption including dysfunction of liver and muscle mitochondria. Tissue-specific molecular signatures, which we have unveiled, clearly indicate that inefficient utilization of glucose is exacerbated by enhanced uptake by the gut. Besides providing systems-wide mechanistic insights, we propose that consumption of SSBs causes intestinal ‘molecular addiction’ for deregulated absorption of hexose-sugars, and drives diseases such as diabetes and obesity.

## Introduction

Sucrose consumption has been proposed to affect physiology and drive the onset of metabolic, immune, and neurocognitive deficits, with obesity and insulin resistance being the most prominent. An alarming increase in sucrose intake has been evidenced across all socio-economic strata, and estimates from the World Health Organization (WHO) indicate that there is a dramatic trend towards overconsumption all over the world [2–6]. While strong correlations have been made between sucrose consumption and the elevated risk of developing obesity, diabetes, and non-alcoholic fatty liver disease (NAFLD) in humans [7–13], causal factors/mechanisms have remained elusive. Given the wide ranging physiological deficits, if and how sucrose overconsumption affects cellular, molecular and metabolic pathways in a tissue specific manner is nearly impossible to address in humans. Furthermore, there is an urgent need to unravel mechanisms that drive or contribute to ill-effects of sucrose over-consumption using preclinical studies. Such studies will be useful to raise new hypotheses to understand and mitigate ill effects of consumption of sucrose in beverages in humans.

Even though several studies in rodents have provided insights, paradigms employed for sucrose consumption have used extremely high concentrations that are rarely observed in humans [14,15]. Moreover, a majority of data has relied on excess sucrose in feed rather than in water, which have been shown to produce distinct and often disparate effects on physiology. It is also important to note that most of our understanding on sucrose overconsumption has emerged from studies that have combined sucrose administration with a high-fat diet or have investigated the ill-effects of overconsumption of fructose [16–29]. However, since increasing evidences have established an alarming rise in consumption of Sugar-Sweetened Beverages (SSBs), there is an urgent need to delineate the molecular and physiological underpinnings of deficits associated with sucrose immoderation. Although few, there are reports that have administered human-relevant levels of sucrose. In this context, seminal insights from recent studies have indicated sex-specific effects [30] of human-relevant sucrose consumption [31]. Notably, while young adult male mice showed a greater increase in body weight gain and higher lean mass, females displayed perturbed tissue-specific lipid homeostasis with no apparent change in body weight. Besides enhanced mortality and reduced fitness [32], [33], sucrose consumption led to de-novo lipogenesis, largely driven by liver-adipose crosstalk [31]. However, a comprehensive analysis of molecular, metabolic, and morphological factors across tissues that drive pathophysiological manifestations of human-relevant sucrose consumption is still lacking. Specifically, if and how mechanisms that dictate nutrient uptake and utilization within organ systems that contribute towards macronutrient metabolism remain to be addressed. Further, since the pathophysiological manifestations have an underlying imbalance between anabolism and catabolism, whether human-relevant sucrose consumption impinges on fed and fasted states differentially remains to be addressed.

Macronutrient metabolism at a system level is inherently dependent upon transport in the gastro-intestinal tract, tissue-specific uptake, and utilization for anabolic and energetic needs. In fact, the small intestine is known to be plastic and undergoes dramatic changes upon changes in dietary composition vis-a-vis villi architecture, cellular responses and inflammation [34–38] . Besides, several reports have indicated biased or skewed uptake and utilization of nutrients in response to high-calorie intake [39,40]. However, whether human relevant sucrose consumption impinges on region-specific intestinal mechanisms that are key for nutrient transport and metabolic sensing remains to be uncovered.

Excess carbohydrate intake is well known to reprogram gene expression, which is also associated with insulin resistance. Current studies employing very high concentrations of sucrose have identified distinct molecular signatures associated with sucrose, fructose and high-fat diet-induced obesogenic phenotypes [41] . Nonetheless, given the interplay between gene expression, insulin signaling and energetics [33,42–45], it is intriguing to find that only a few studies have attempted to tease out multi-organ system responses following dietary perturbations, which are close to real-life scenarios. Specifically, mitochondrial functions across tissues that contribute to systemic metabolism have not been addressed following human-relevant sucrose consumption. This is important since, along with intestinal uptake, capturing tissue-specific molecular and physiological parameters will be key to provide a cause-consequence relationship.

In this study, we have elucidated metabolic, molecular, cellular, and mitochondrial underpinnings of excess nutrient uptake as a consequence of chronic consumption of 10% sucrose in water, which is relevant owing to the fact that commonly consumed beverages have 10-15% sucrose. Our results provide hitherto unknown tissue-specific responses and highlight that moderate sucrose consumption elicits distinct responses in the intestine, liver, and muscles, which are associated with insulin resistance and obesity.

## Materials and Methods

### Mice experiments

C57BL/6 mice housed under standard animal-house conditions were used for experiments. The procedures and the project were approved and in accordance with the institute animal ethics committee (IAEC) guidelines. 2.5-3-month old male mice were either administered normal water (CW) or 10% sucrose in drinking water (SW) ad libitum, along with normal chow feed for a period of three months. Mice were regularly observed for infections and appropriate measures were taken to maintain the general health of the mice.

After 3 months of Sucrose feeding, the mice were either starved overnight before sacrifice at 9.30am and sample collection, or they were fed ad-libitum followed by I.P glucose injection at 9am and sacrificed at 9.30am.

### Body weight Measurements

Body weight measurements were taken weekly at 10.00AM using an analytical weighing balance during the three month dietary perturbation.

### Blood Glucose measurements

Blood glucose levels were monitored as indicated from tail vein prick using AccuCheck Glucometer, Roche.

### Tolerance Tests

#### Pyruvate and Insulin Tolerance Test (PTT and ITT)

Mice were fasted for 6h (with normal drinking water), after which body weight and blood glucose levels were measured. 2g/Kg b.wt of pyruvate in PBS (for PTT) or Insulin (0.65 IU/kg b.wt, HumulinR, Lilly) (for ITT) was injected into each mouse intraperitoneally. The blood glucose level was monitored with a glucometer at indicated intervals during a 2h time course. Area under the curve (AUC) was plotted using GraphPad Prism version 8.

#### Intraperitoneal Glucose Tolerance Test (IP-GTT)

Mice were fasted for 12-14h (with normal drinking water), after which body weight and blood glucose levels were measured. 2g/Kg b.wt of glucose was injected into each mouse intraperitoneally. The blood glucose level was measured with a glucometer at indicated intervals during a 2h time course. Area under the curve (AUC) was plotted using GraphPad Prism version 8.

#### Immunofluorescence and histology

Mice were sacrificed by cervical dislocation and tissue segments from all three regions of the intestine were isolated, flushed with PBS and fixed in 4% PFA in PBS overnight at 4°C, and then either embedded in Technovit 7100 resin (Electron Microscopy Sciences, #14653) for histological examination or processed further for immunofluorescence staining.

Fixed samples were serially dehydrated with ethanol in PBS (30%, 50%, 70%, 90%, 99%), and embedded in Technovit 7100 resin. The embedded blocks were sectioned using a microtome (Leica RM2265) to obtain 3µm thick transverse sections which were then stained with 0.4% toluidine blue or Giemsa for histological studies.

### Immunofluorescence staining for crypt proliferation

Fixed samples were postfixed in 100% methanol and then stored at -20°C. Prior to staining, the outermost muscularis was manually removed under a bright field microscope. For permeabilization, tissues were briefly treated with Proteinase K (Sigma) and then washed in PBT (0.8%, Triton X-100 in PBS). Samples were blocked with 10% NGS for four hours at room temperature, and then incubated with rabbit anti-Ki67 (1:200; Invitrogen, MA5-14520) primary antibody overnight at room temperature. Primary antibody was then washed off with PBT and treated with anti-rabbit secondary antibody at room temperature for six hours. Secondary antibody was then washed off and samples were postfixed in 4% PFA in PBS overnight at 4°C. The secondary antibody solution used included Alexa 488 conjugated anti-rabbit antibody (1:200; Invitrogen, A11034), and DAPI (1:200).

Samples were then embedded in Technovit 7100 resin after dehydration with ethanol in PBS as mentioned in the above section, and 6µm thick transverse sections were taken using the microtome. Sections were mounted in Fluroshield mounting medium (Sigma) before imaging.

### Image acquisition and quantification

#### Histology

The samples were imaged on a stereoscope, SteREO Discovery (Zeiss) using AxioCam (Zeiss). Eighteen consecutive sections were taken for each animal. Of these, five random sections were imaged for each animal. ImageJ software was used to measure the changes in villus morphology. Segmented lines were drawn manually down the centre of the villi for the measurement of the villus height (distance from the villus tip to the base of the villus) and epithelium height (distance from villus tip to the bottom of the crypts). Villus width was measured along the base of the villus, right above the crypts. For each animal, five intact villi were selected at random from the sections. The length of a given villus was measured in all five sections, and the greatest measured length was taken into consideration as the true height. This section was assumed to pass through the centre of the villus, and measures for the epithelium height and villus width were taken from this section. Intestines from total 8 overnight starved animals from 4 cohorts for each condition were used to quantify villi characteristics mentioned above for duodenum in males. For Jejunum and ileum, 9 and 7 control, and 10 and 8 sucrose fed animals, were quantified, respectively. In addition, two overnight starved female cohorts, consisting of 6 control and 5 sucrose fed animals for duodenum, 5 each for jejunum, and 7 controls and 5 sucrose fed animals for ileum were quantified.

#### Immunofluorescence staining

Tissue sections were imaged using Zeiss 880 confocal microscope and at least three sections were imaged for each animal using Plan-APOCHROMAT 40×/1.3 lens. Six crypts were selected at random and cells showing DAPI and Ki67 signals were counted manually using the ImageJ Cell Counter plugin. The number of Ki67 positive nuclei and DAPI labelled nuclei were summed up from the 6 crypts for each animal. The ratio of the number of crypt cells positive for Ki67 and the total number of cells as stained by DAPI was used as a readout for cell proliferation for each animal.

#### Goblet cell counts

3μm thick tissue sections were stained with Giemsa stain for 15 minutes on a heating block, followed by destaining with 0.1% acetic acid for 10 seconds. The sections were then imaged on a Zeiss Apotome fitted with an AxioCam MRc5 using a 10×/0.3 Plan-NEOFLUAR lens. Two consecutive sections were imaged for each animal. The tissue area in each section was measured, and goblet cells were counted using the ImageJ Cell Counter plugin. Tissue area and goblet cell numbers from both sections were averaged, and goblet cell numbers/area was determined for each animal, for each segment of the intestine. Goblet cell numbers were determined for two cohorts of fed male mice. Goblet cell numbers were counted from the duodenum of 8 sucrose fed and control animals each while for the jejunum of 10 animals for each condition were used. For ileum, 10 control animals and 9 sucrose fed animals were used for the goblet cell counts.

### Mitochondrial isolation

Liver and quadriceps muscle were excised and minced in ice-cold Mitochondrial Isolation Buffer (70mM Sucrose, 220mM Mannitol, 10mM HEPES, 1mM EGTA, 2mg/ml BSA, pH 7.2). Minced tissue was homogenized using a motor-driven Teflon-glass homogenizer (2-3 strokes for liver, 15-18 strokes for muscle), following which the homogenate was spun at 800xg, 10min, 4c twice to pellet cellular debris and nuclei. The supernatant was collected and spun at 12000xg, 10min, 4c twice and the final pellet was resuspended in Mitochondrial Assay Solution (70mM Sucrose, 220mM Mannitol, 10mM KH_2_PO_4_, 2mM HEPES, 1mM EGTA, 5mM MgCl_2_, 4mg/ml BSA, pH 7.2).

### OCR and ATP measurements

OCR was measured using a Seahorse Xfe24 Analyzer. Immediately post isolation, mitochondria were plated in a Seahorse plate at 3-10ug/50ul in each well, depending on the substrate used for respiration. Complex-specific substrates were used to profile respiration - Pyruvate/Malate (7.5mM/2.5mM) were used for complex 1, Succinate/Rotenone (10mM/2uM) were used for complex 2, and Palmitoylcarnitine/Malate (50uM/5mM) was used for FAO-dependent respiration. The plate was centrifuged at 2000xg, 20min, 4c to ensure uniform adherence of mitochondria, after which MAS containing specific substrates were added in each well. The plate was incubated in a non-CO2 incubator at 37°C for 8mins, after which OCR was measured using sequential injections of ADP (100uM), Oligomycin (4uM), FCCP (4uM), Rotenone/Antimycin A (2uM/4uM). In parallel, mitochondria were incubated with the same substrates and ADP in 1.5ml microcentrifuge tubes at 37°C for 30 mins, after which the reaction mixture was used to measure ATP levels via a luciferase/luciferin based assay system according to manufacturer’s protocol (FLAA - Sigma Aldrich).

### ROS production assay

Mitochondria were isolated as per protocol and diluted to 5ug/12.5ul in MAS buffer. 25ul of 4X substrate stock was added to each well of a 96-well black plate. AR stock was diluted to 400uM and an equal volume of this dilution was added to the mitochondrial suspension. 25ul of this Mitochondria-AR mixture was added to each well. Next, 50ul of a 20U/ml HRP solution was added to each well to start the reaction. Excitation/emission wavelengths were set at 530/590nm and fluorescence was measured every 5min for 1hr or until the standards saturated.

*Standards:* 3% H2O2 was diluted to a 20mM stock. From this, a mixture of 10uM H2O2 and 100uM AR was made and 0,10,20,30,40,50ul of this mixture was added to wells in duplicates. The volume was made to 50ul with MAS buffer.

*Solutions:*

● 10mM Amplex Red (AR) in DMSO
● 10mg/ml Horseradish Peroxidase (HRP) in 0.1M Potassium Phosphate Buffer, pH 6.0

#### RNA Isolation and qRT-PCR

TriZol reagent was used for the isolation of total RNA from liver, small intestine and skeletal muscle samples. 1ug of total RNA was used to prepare cDNA using Random Hexamers and SuperScript IV Reverse Transcriptase. The sample cDNA was used for qRT-PCR using KAPA SYBR FAST qPCR Master Mix (2X) Kit and LightCycler 96/480 (Roche) Instrument. The protocols prescribed by manufacturers were followed for RNA isolation, cDNA preparation and qRT-PCR. Expressions of genes of interest were normalized to that of Actin (Actb)/18s rRNA transcripts. Data was plotted using GraphPad version 8, and t test was used for statistical analysis.

#### Lysate prep and western blotting

Isolated liver and small intestinal samples were minced in ice-cold RIPA buffer(Tris-Cl 10mM pH 7.4, NaCl 140mM, Sod. deoxycholate 0.1%, SDS 0.1%) supplemented with Protease Inhibitor Cocktail, PhosSTOP and PMSF and incubated for 15 minutes in ice. The homogenate was spun at 12,000rpm at 4°C for 15 mins, following which the supernatant was collected and to it 4X Laemmli was added to a final concentration of 1X.

GS muscle was excised and minced in ice-cold RIPA buffer(Tris-Cl 10mM pH 7.4, NaCl 140mM, Sod. deoxycholate 0.1%, SDS 0.1%) supplemented with Protease Inhibitor Cocktail, PhosSTOP and PMSF and incubated for 40 minutes in ice with intermittent vortexing. The homogenate was spun at 12,000rpm at 4C for 15 mins, following which the supernatant was collected and to it 4X Laemmli was added to a final concentration of 1X.

Protein estimation was done using BCA assay, and 30ug of total protein was loaded into each lane of an SDS-PAGE gel, followed by western blotting for indicated proteins. Protein bands were then quantified against either total protein levels obtained from Ponceau Staining, or against loading control as mentioned. Data was plotted in GraphPad version 8, and t test was used for statistical analysis.

#### Apical membrane fraction isolation and western blotting

Apical membrane fraction was isolated from frozen intestine samples based on the protocol proposed by [1]. ∼80-100 mg of frozen intestinal sample was crushed in 500ul of ice cold Tris-Mannitol buffer (pH=7.1) supplemented with CaCl_2_ (final concentration of 10mM) and incubated in ice for 15 minutes with intermittent vortexing. The homogenate was centrifuged at 4°C for 10 minutes at 2000g. The supernatant was collected and spun at 4°C for 15 minutes at 20,000g. The yellowish brown pellet was the desired membrane fraction which was dislodged in 120ul Tris-mannitol buffer and to it 4X Laemmli was added to a final concentration of 1X.

Protein estimation was done using BCA assay, and 50ug of total protein was loaded into each lane of an SDS-PAGE gel, followed by western blotting for indicated transporter protein. Protein bands were then quantified against total protein levels obtained from Ponceau Staining. Data was plotted in GraphPad version 8, and t test was used for statistical analysis.

#### Sucrose hydrolysis assay

Frozen intestinal tissues were homogenized in Phosphate Buffer (pH=6.8) supplemented with PIC (Roche) at room temperature and centrifuged at 3000g, 4°C for 10 minutes. The supernatant was collected and total protein was estimated. 1:10 dilutions were made for the supernatants and incubated with equal volumes of 25mM Sucrose solution at 37°C for 15 minutes. Glucose concentration for the final reaction mix was estimated using Glucose Colorimetric Detection Kit (Invitrogen) as per the manufacturer’s protocol.

**List of primers used in this paper:**

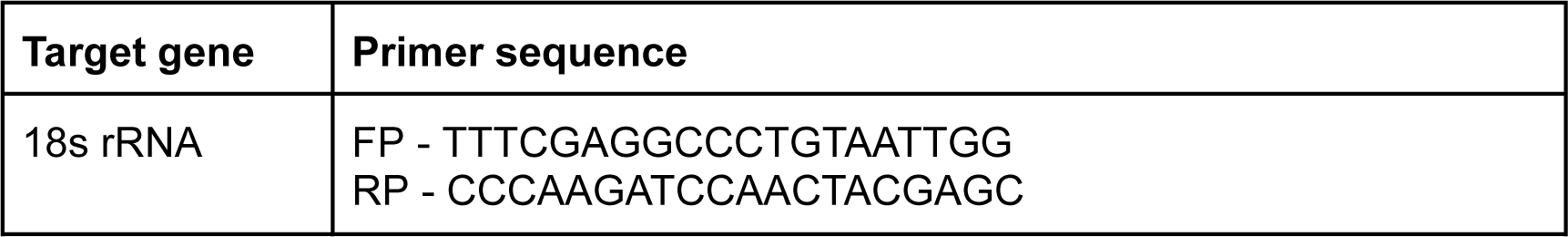
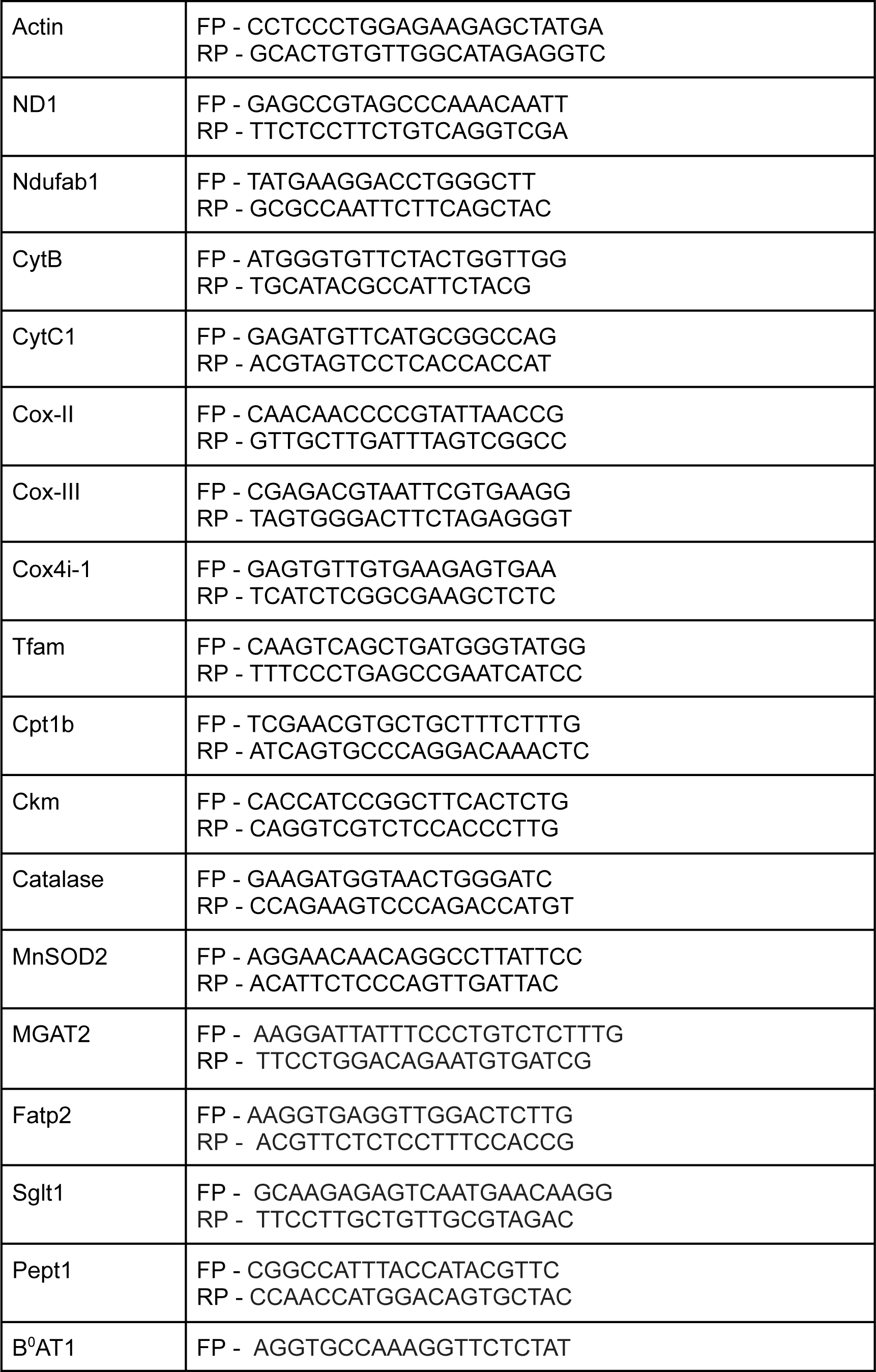
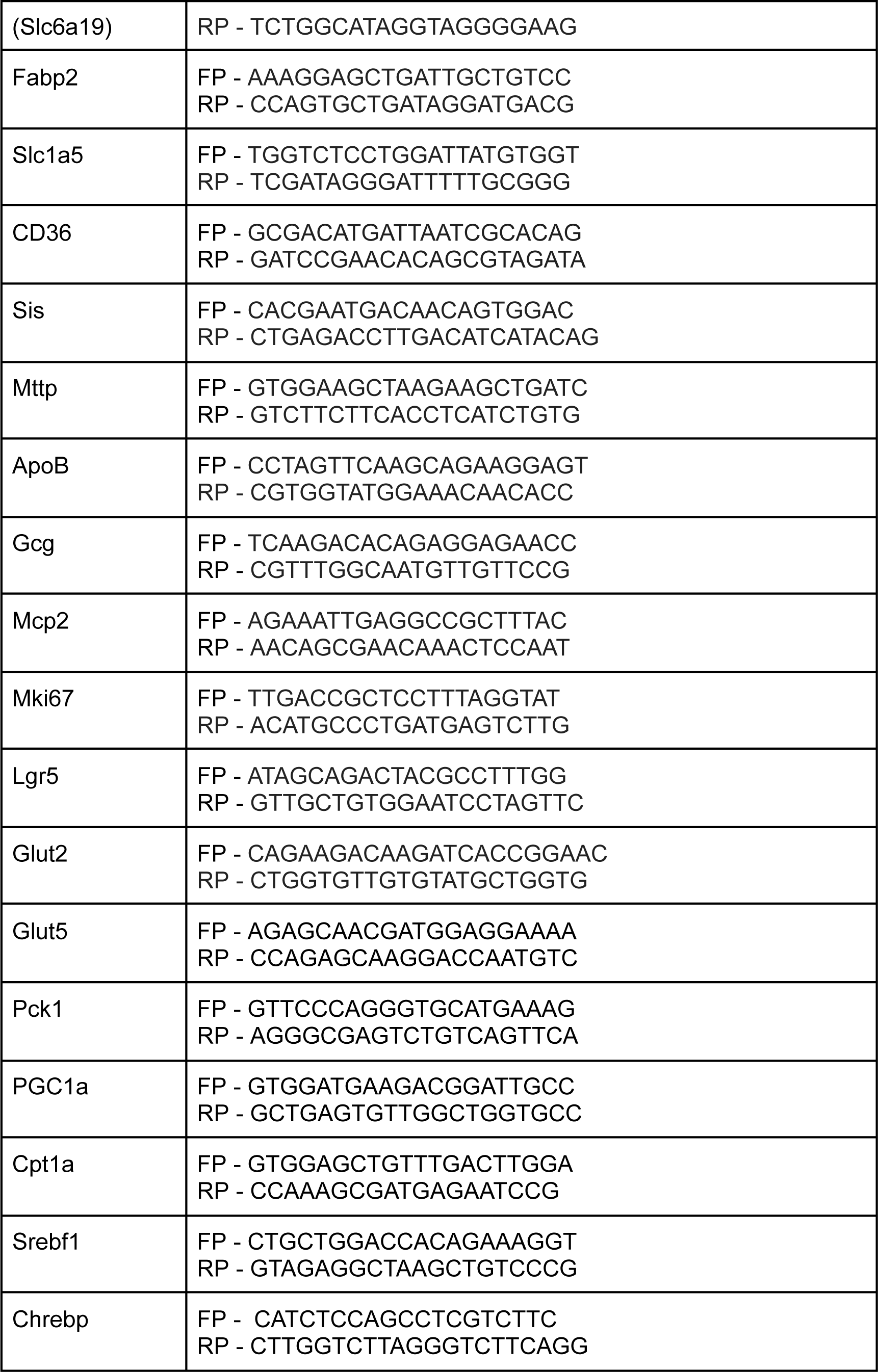
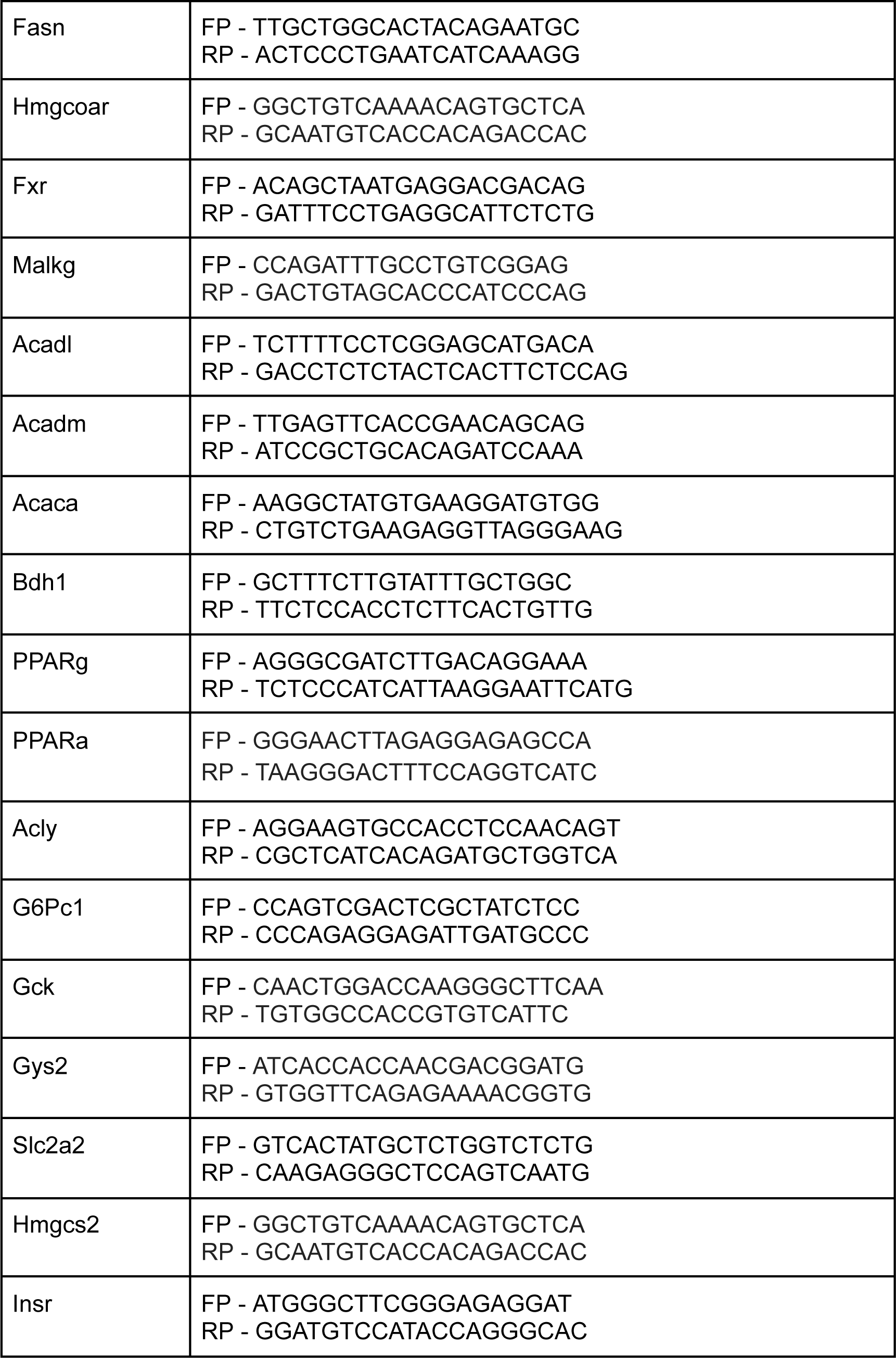
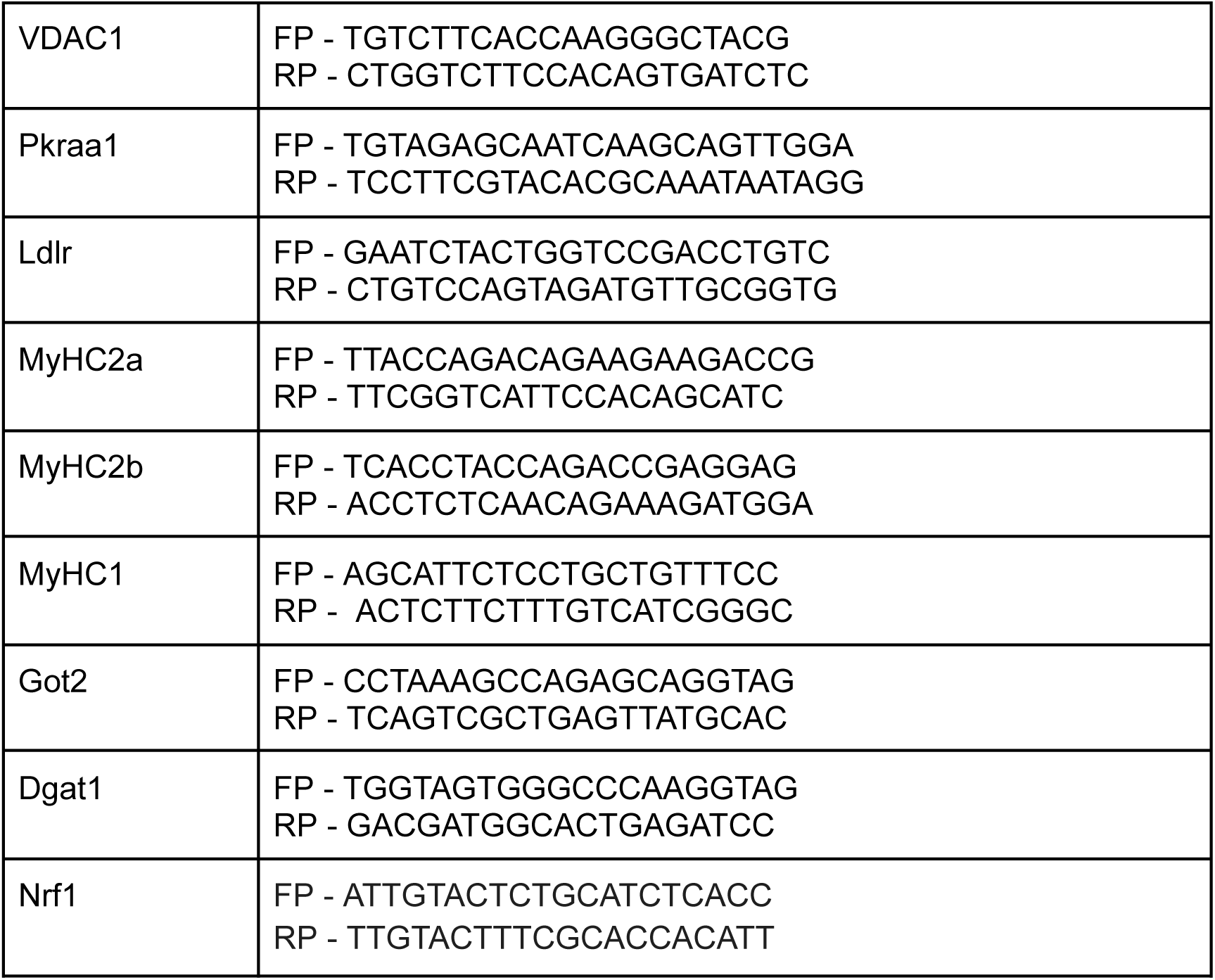

**List of reagents used in this paper:**

**List of reagents used in this paper :**
| Reagent | Source | Cat. No. |
| --- | --- | --- |
| Sucrose | Himedia | MB025 |
| D-Mannitol | Merck | 443907 |
| HEPES | USB | 16926 |
| EGTA | Sigma | E3889 |
| MgCl<sub>2</sub> | USB | 18641 |
| CaCl<sub>2</sub> |  |  |
| MgSO<sub>4</sub> |  |  |
| BSA | Sigma | A7030 |
| Protease Inhibitor Cocktail | Sigma | 5056489001 |
| PhosSTOP | Sigma | 4906837001 |
| PMSF | Sigma | P7626 |
| KH<sub>2</sub>PO<sub>4</sub> | Sigma | 60218 |
| D+Glucose | Sigma | G7021 |
| Sodium Pyruvate | Sigma | P2256 |
| Insulin | Lilly |  |
| BCA | Sigma | B9643 |
| Sodium Succinate | Sigma | 14160 |
| L-Malic Acid | Sigma | M6413 |
| Palmitoyl-L-Carnitine Chloride | Sigma | P1645 |
| ADP | Sigma | A2754 |
| ATP | Sigma | A2383 |
| Rotenone | Sigma | R8875 |
| Oligomycin A | Sigma | 75351 |
| FCCP | Sigma | C2920 |
| Antimycin A | Sigma | A8674 |
| Cytochrome C | Sigma | C2506 |
| Luciferin-Luciferase Mix | Sigma | FLAAM |
| PVDF Membrane | Immobilon | IPVH00010 |
| Trizol Reagent | Ambion | 15596018 |
| Chloroform | Qualigens | Q12305 |
| Isopropanol | Fisher Bioreagents | BP2618-212 |
| DEPC treated water | Ambion | AM9915G |
| SSIV-RT, SSIV Assay Buffer,<br>DTT | Invitrogen | 18090050 |
| dNTP | Genei | 065100003(A-D) |
| RNaseOUT | Invitrogen | 10777019 |
| Random Hexamers | Invitrogen | N8080127 |
| KAPA SYBR FAST qPCR Master Mix (2x) | KAPA Biosystems | KK4602 |
| Glucose Colorimetric Detection Kit | Invitrogen | EIAGLUC |
| Amplex UltraRed | ThermoFisher Scientific | A12222 |
| Peroxidase Type VI from Horseradish | Sigma-Aldrich | P8375 |
| 3% H2O2 | Sigma-Aldrich | 88597 |

## Results

### Sucrose feeding in water induces glucose intolerance

We set out to assess the physiological and molecular effects that emerge as a consequence of sucrose overfeeding. Towards this, adult mice reared on normal chow diet were provided ad-libitum access to 10% (w/v) sucrose in water for 3 months and mice that had access to water alone acted as controls (Figure 1A). We specifically chose C57BL6/N strain, unlike the prevalent literature on C57BL6/J, because the latter harbors a mutation in the *Nnt* gene and is known to affect metabolic responses including to dietary perturbations. In addition to euthanizing under ad-libitum fed state, both groups of mice were starved overnight to score for fasting responses, as described below (Figure 1A). We observed a significant increase in body size and weight in sucrose-fed mice [31,39] when compared to controls (Figures 1B,1C). While intriguing, it is unclear if the increase in the consumption of 10% sucrose water and the associated reduction in total food intake (Figure 1D, 1E) was consequential or compensatory. The whole-body energetics, as assessed by respiratory exchange ratio (RER), was not different between the control and sucrose water fed animals (Figure S1A). Importantly, there was a consistent elevation in circulating glucose levels (Figure 1F), which was independent of the time of the day, indicating hyperglycemia, consistent with a recent report [31]. The mild weight gain observed in females in our study could possibly be strain dependent (Figure S1B).

**Figure 1:**
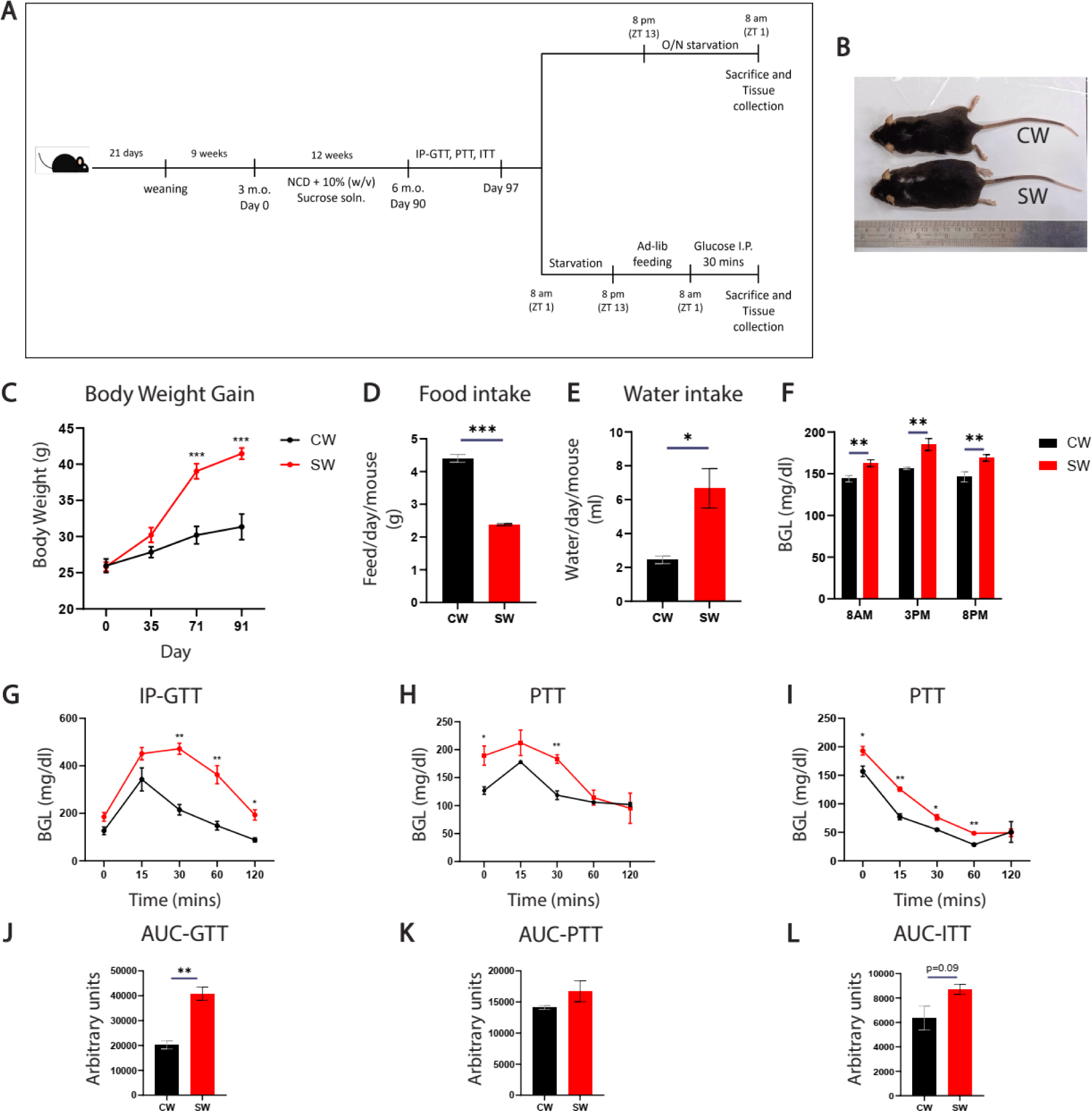
10% Sucrose feeding in water induces glucose intolerance. (A) Schematic representation of experimental paradigm used to study the effects of 10% Sucrose overfeeding for 3 months in C57B/6N male mice. (B) Representative image of mice fed with NCD + Water (CW) and NCD + 10% Sucrose soln. (SW). (C) Total body weights of mice, (D) intake of food (E) and water intake in CW and SW mice. (F) Blood Glucose levels measured at 8am, 3pm, 8pm in ad-lib fed CW and SW mice . Blood glucose levels for (G) IP-GTT in overnight fasted mice, (H) PTT and (I) ITT carried out on 6h fasted mice (N=4, n=4-6). (J-L) AUC quantifications for blood glucose levels observed during (J) IP-GTT (K) PTT and (L) ITT. Experimental and technical repeats; N=4, n=4-6 for all experiments. Data represented as Mean ± SEM and analysed by t-test. P-value of 0.05 was considered significant. *P ≤ 0.05; **P ≤ 0.01; ***P ≤ 0.001.

To assess the underlying physiological basis of hyperglycemia, which has not been addressed earlier, glucose-, insulin- and pyruvate-tolerance tests (GTT, ITT & PTT, respectively) were performed. As shown in Figures 1G-1I, sucrose-fed mice displayed impaired glucose homeostasis with significantly elevated responses on all three measures. Specifically, sucrose fed mice were unable to clear exogenously administered glucose at all time points measured during GTT (Figures 1G, 1J and Figure S1E). We also observed perturbed response in sucrose-fed mice during PTT (Figures 1H, 1K and Figure S1F) and ITT (Figures 1I, 1L), which indicated that impaired glucose homeostasis could be cumulatively contributed by both increased hepatic gluconeogenesis and reduced insulin signaling. Unlike the differences in the weight gain, we found both males and females had similar impairment in glucose homeostasis.

### Histological remodeling of the intestine

Over or under-nutrition, and paradigms of excess dietary carbohydrate/fat have been associated with histopathological changes, which are both causal and consequential vis-a-vis impaired metabolism. Earlier studies, which have largely employed either very high concentration of sucrose water or have combined it with high-fat diets, have reported such changes mostly in the liver [29,31,39,45,46]. Specifically, to score for alterations in tissues that govern nutrient absorption and central metabolic homeostasis, we assessed histology of the muscle, and the small intestine. SDH-based staining and molecular assays showed that proportions of oxidative and glycolytic fibres were similar (Figure S2H) and there were no indications of atrophy or hypertrophy of muscle following 10% sucrose-water over-feeding.

However, it was interesting to note that there was a subtle but statistically significant increase in the heights of duodenal and jejunal villi in sucrose fed male mice when compared to the controls (Figures 2A, 2B). This change was also apparent in the duodenum and jejunum of female mice fed with sucrose (Figure S2A, S2B). This was in agreement with previous literature where prolonged consumption of high caloric diets increased the absorptive surface area of the small intestinal epithelium, thereby predisposing organisms to overnutrition [34,38]. These data clearly suggested that the small intestine was prone to morphological remodeling in response to 10% sucrose-water consumption, the muscles.

**Figure 2:**
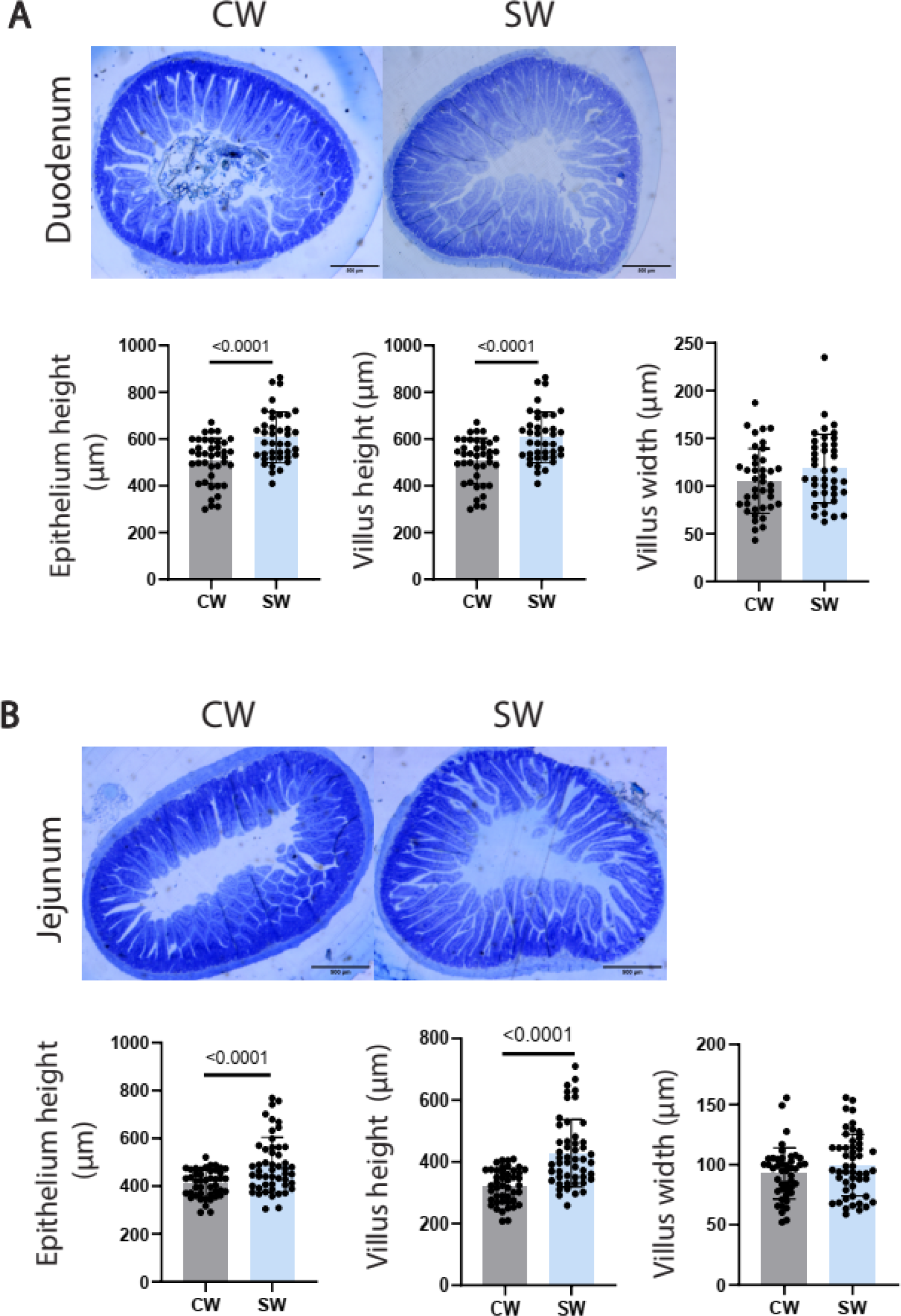
Histological remodeling of the Small Intestine. (A,B) Representative histological sections and quantification of morphological parameters of (A) Duodenum and (B) Jejunum displaying the impact of sucrose overfeeding on epithelial morphology. Sections obtained from CW and SW mice following overnight fasting. Data represented as Mean ± SD and analysed by t-test (N=7-10, n=40-50). CW - NCD + Water, SW - NCD + 10% Sucrose soln. P-value of 0.05 was considered significant. *P ≤ 0.05; **P ≤ 0.01; ***P ≤ 0.001.

### Tissue-specific rewiring of gene expression in sucrose overfed mice under overnight fasted and fed states

Others and we have shown earlier that allostatic load associated with diet-induced physiological changes could emerge as a consequence of perturbed expression of metabolic genes and/or metabolic alterations independent of molecular rewiring [47–53]. Based on the physiological changes described above, we wanted to measure transcript levels of genes involved in glucose and fat metabolism, transcriptional regulation and mitochondrial functions across tissues. More importantly, we investigated the molecular signatures under overnight fasted and ad-libitum fed conditions in both control and 10% sucrose fed cohorts, also to correlate with anabolic and catabolic drives in these states, which is hitherto unknown.

It was interesting to note that the mRNA expression of Pck, G6Pc, Lcad, Mcad, Sirt1, Ppara, Ppargc1a, Chrebp, Dgat1, Pparg and others was similar in the liver between control and sucrose cohorts as indicated (Figures 3A, 3B, 3C, S4A, S4B; Figures 4A, 4B, S5C). This was consistent with no change in protein levels of key factors such as SIRT1 (Figure S3A, S5A), the upstream regulator of metabolic genes. However, we found PPARGC1A (PGC1A) (Figures 3D, 4C) and FOXO1 (Figures 3E, 4D) proteins to be reduced in 10% sucrose water fed mice in ad-libitum fed and fasted conditions. This posited a gene-expression independent metabolic rewiring in the liver, especially for enhanced hepatic gluconeogenesis. While there was a reduced expression of lipogenic genes in response to sucrose consumption (as indicated in Figure 4B), transcripts levels of CD36, one of the fatty acid transporters, was significantly higher (Figures 3C, 4B), indicating perturbed hepatic lipid homeostasis. Moreover, unlike the obesogenic diets that affect skeletal muscle lipid metabolism, [45,54], we found no change in genes involved in fatty acid metabolism following sucrose water consumption (Figure 3F).

**Figure 3:**
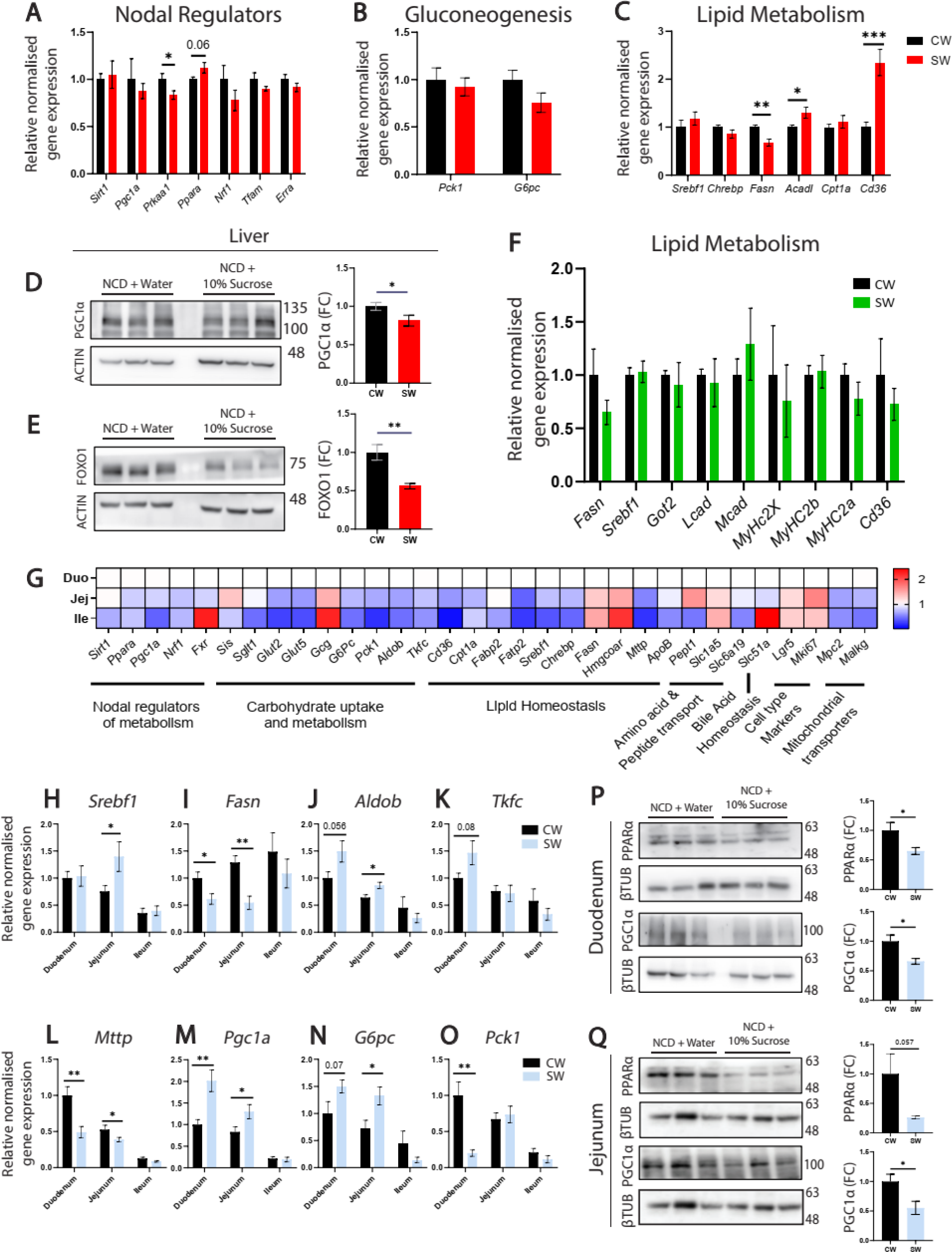
Tissue-specific molecular rewiring under a starved state. (A-C) Relative hepatic gene expression analysis of CW and SW mice under 12hr/overnight fasted conditions. (A) Nodal regulators of metabolic responses, (B) gluconeogenic genes and (C) genes regulating lipid metabolism, as indicated . (D,E) Representative immunoblot for (D) Pgc1α and (E) FOXO1 from lysates of liver from 12hr/overnight starved CW and SW mice. (F) Relative expression of genes regulating lipid metabolism in gastrocnemius of 12hr/overnight fasted CW and SW mice. (G) Relative intestinal (Duodenum, Jejunum and Ileum) gene expression analysis of CW mice under 12hr/overnight fasted condition as indicated. Data represented as fold change with respect to baseline expression in the duodenum. (H-O) Relative intestinal (Duodenum, Jejunum and Ileum) gene expression analysis of CW and SW mice under 12hr/overnight fasted condition. (H,I) De-novo lipogenesis (J,K) fructose metabolism (L) LD synthesis (M,N,O) gluconeogenesis and β-oxidation . Data represented as fold change with respect to expression in CW Duodenum. (P,Q) Representative immunoblot for PPARα and Pgc1α in (P) Duodenum and (Q) Jejunum of CW and SW fed mice under 12hr/overnight fasted condition. CW - NCD + Water, SW - NCD + 10% Sucrose soln. Experimental and technical repeats; N=2-3, n=3. Data represented as Mean ± SEM and analysed by t-test. P-value of 0.05 was considered significant. *P ≤ 0.05; **P ≤ 0.01; ***P ≤ 0.001.

**Figure 4:**
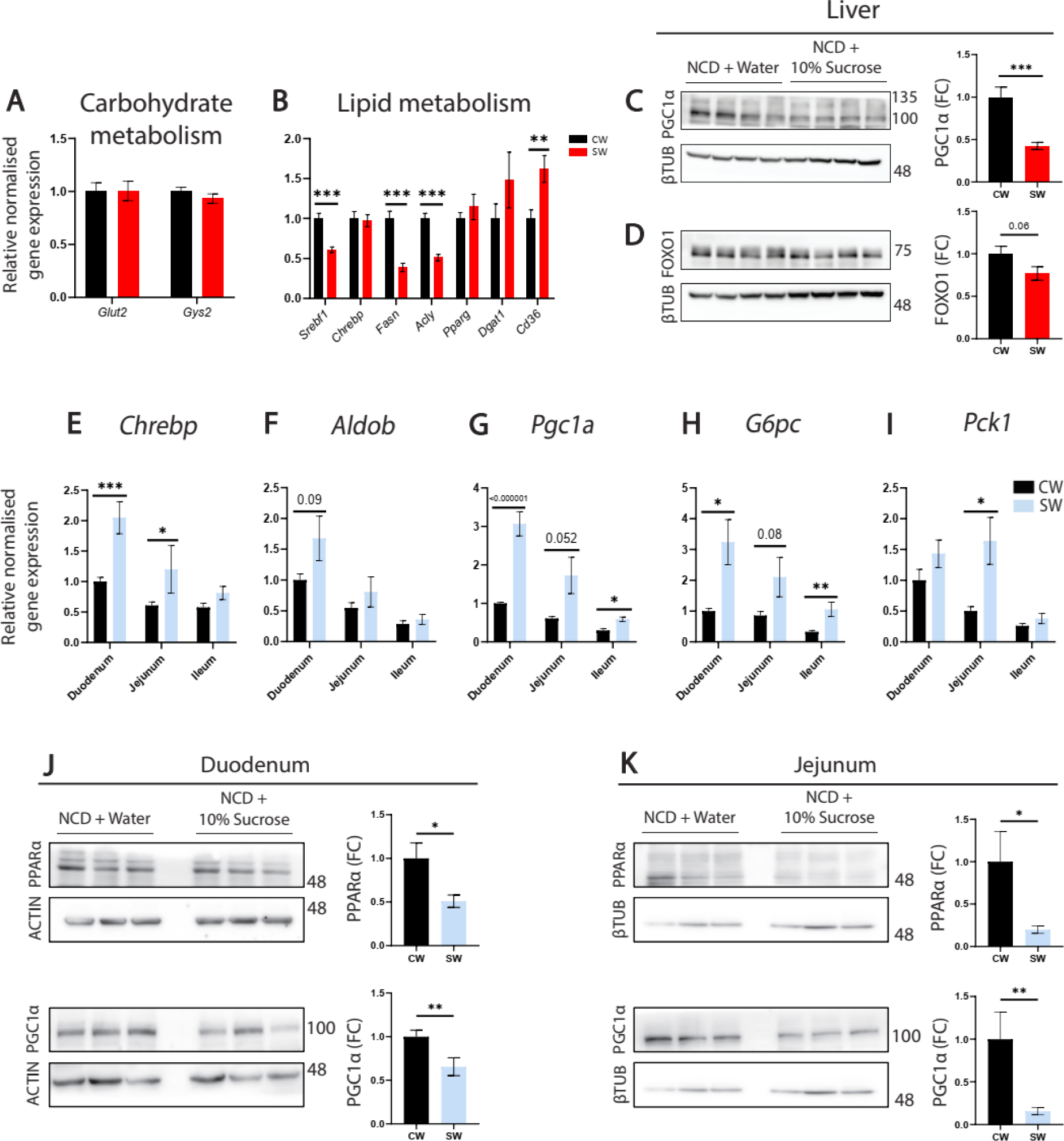
Tissue-specific molecular rewiring under a fed state. (A,B) Relative hepatic gene expression analysis for (A) Glucose transporter (Glut2) and glycogen Synthase (Gys2) and (B) lipid metabolism in CW and SW mice as indicated. (C,D) Representative immunoblot for (C) Pgc1α and (D) FOXO1 in the lysates of liver from CW and SW mice, as indicated. (E-I) Relative intestinal (Duodenum, Jejunum and Ileum) gene expression analysis of (E) Chrebp (de-novo lipogenesis and fructose metabolism) (F) Aldob (fructose metabolism) (G) Pgc1a (gluconeogenesis and β-oxidation) (H,I) gluconeogenic genes (G6Pc and Pck1) from CW and SW mice, as indicated. Data represented as fold change with respect to expression in CW Duodenum. (J,K) Representative immunoblot for PPARα and Pgc1a in the lysates of (J) Duodenum and (K) Jejunum from CW and SW mice. Experimental and technical repeats; N=2, n=3-4. Intraperitoneal glucose was administered to the ad-libitum fed mice to assess insulin responsive molecular changes in a ‘postprandial state’. Data represented as Mean ± SEM and analysed by t-test. P-value of 0.05 was considered significant. *P ≤ 0.05; **P ≤ 0.01; ***P ≤ 0.001.

Although the small intestine plays a deterministic role in regulating systemic physiology owing to its ability to transport nutrients, it is surprising that relatively less is known about gene expression changes as a function of dietary alterations in this tissue, when compared to liver, muscle and adipose tissues. This is further complicated because of overlapping and non-overlapping functions of duodenum, jejunum and ileum [55] and nutrient/metabolic gradients along both the length of the gut and across apico-/basal-surfaces, and under fed and fasted conditions. Thus to characterize baseline expression, we performed real-time PCR analyses in response to 12-hours starvation in control cohorts and found that region specific expression of genes such as Sglt1, Gcg, Slc51a, Mttp and FxR were consistent with earlier reports [56–58]. Levels of transcripts associated with nutrient uptake and utilization including glucose and fat metabolism, bile-acid homeostasis and transcriptional regulators also displayed distinct patterns of expression, in accordance with known functions of duodenum (major site of dietary carbohydrate and fatty acid uptake), jejunum (lipid droplet packaging and chylomicron secretion) and ileum (bile acid reuptake, short-chain fatty acid uptake, GLP1 release) (Figure 3G).

Next, we investigated the impact of 10% sucrose overfeeding on the small intestine, under both ad-libitum fed and fasting conditions. In addition to taking care of the feeding dependent variability, sampling following 12-hours of fasting was necessary to characterize the molecular responses of duodenum, jejunum and ileum under a fasted state, which is less explored (and as a consequence of chronic consumption of sucrose water).

It was interesting to note that transcriptional regulators involved in fat and glucose homeostasis showed altered expression in a region-dependent manner in the sucrose cohort under both fed and fasted states (Figures 3H-O, 4E-I). For example, while Ppargc1a (Figures 3M, 4G) mRNA was increased in sucrose cohorts in *ad libitum* fed and fasted conditions, Srebf1 was increased only in a fasted state (Figure 3H, S5H) and Chrebp expression was increased only in a fed state (Figures 4E, S4C). There was no alteration in SIRT1 protein levels across the regions (Figure S3H-J, S5F-G). Strikingly, levels of the nodal transcription factor PPARa and its coactivator PPARGC1a (PGC1a) proteins, that respond to metabolic cues, were significantly lower in both duodenum and jejunum in the sucrose cohort in both fed and fasted states (Figures 3P-Q, 4J-K). It is important to note that both these factors are known to exhibit tissue-specific expression of alternatively spliced variants [59–61] and our result does not rule out protein level changes of specific isoforms.

To further delineate expression of genes involved in intestinal physiology, specifically in the context of sucrose consumption, we assessed the expression levels of AldoB, Tkfc, G6Pase, Pck1, Fasn and Mttp, key regulators of fructose, glucose and lipid metabolism. We found a robust upregulation of G6Pase in the duodenum and jejunum in sucrose cohorts irrespective of whether the mice were fed or fasted (Figures 3N, 4H). Notably, AldoB and Tkfc (rate limiting enzymes for fructose utilization), and Mttp (rate limiting for pre-chylomicron synthesis) showed a significant change upon sucrose feeding. In the duodenum and jejunum of sucrose fed mice, Aldob showed trends toward increased expression both in the fed and overnight starved conditions (Figures 3J, 4F). At the same time, duodenum of sucrose fed mice showed higher Tkfc expression (Figure 3K) and lower Mttp expression (Figure 3L) in duodenum and jejunum in response to starvation.

To conclude, our results clearly define tissue specific gene expression changes that are associated with 10% sucrose water consumption and indicate the intestine to be more prone to molecular alterations than central metabolic organs like the liver and muscle.

### Differential regulation of nutrient and metabolic sensing/signaling upon sucrose overfeeding

Nutrient sensing and metabolic signaling cascades, that respond to both acute and chronic perturbations, are well established to drive physiological changes via mechanisms including gene expression, transporter expression/localization and modulation of metabolic enzymes [62–69]. Given this and the impaired glucose homeostasis observed in sucrose fed mice (Figure 1), we next assayed for nodal nutrient-sensing pathways viz. AMPK, mTOR and insulin signaling. While AKT is central to insulin signaling, AMPK and TOR kinases sense AMP/ADP/ATP ratios and amino acids, respectively, besides integrating signaling from diverse extracellular cues. As detailed earlier, in order to capture trans-tissue effects that unveil baseline and nutrient induced alterations in metabolic sensing, we assayed for key phosphorylation events in response to intraperitoneal glucose administration under an ad-libitum fed state and following 12-hours fasting in the liver, muscle and intestine (duodenum and jejunum).

Hepatic S473-phosphorylation of AKT (pS473-AKT) was significantly lower following intraperitoneal administration of glucose in ad-lib fed mice (Figure 5A) and showed a trend towards decrease in response to 12-hour starvation (Figure 5B). This clearly indicated that reduction in insulin signaling was more pronounced in the fed state than upon fasting and corroborated impaired glucose homeostasis and insulin resistance observed in the sucrose cohorts (Figure 1). Interestingly, while TOR signaling was significantly downregulated (Figure 5C), AMPK signaling showed a non-significant decrease (Figures S3B, S5B) in the sucrose fed mice livers when compared to controls.

**Figure 5:**
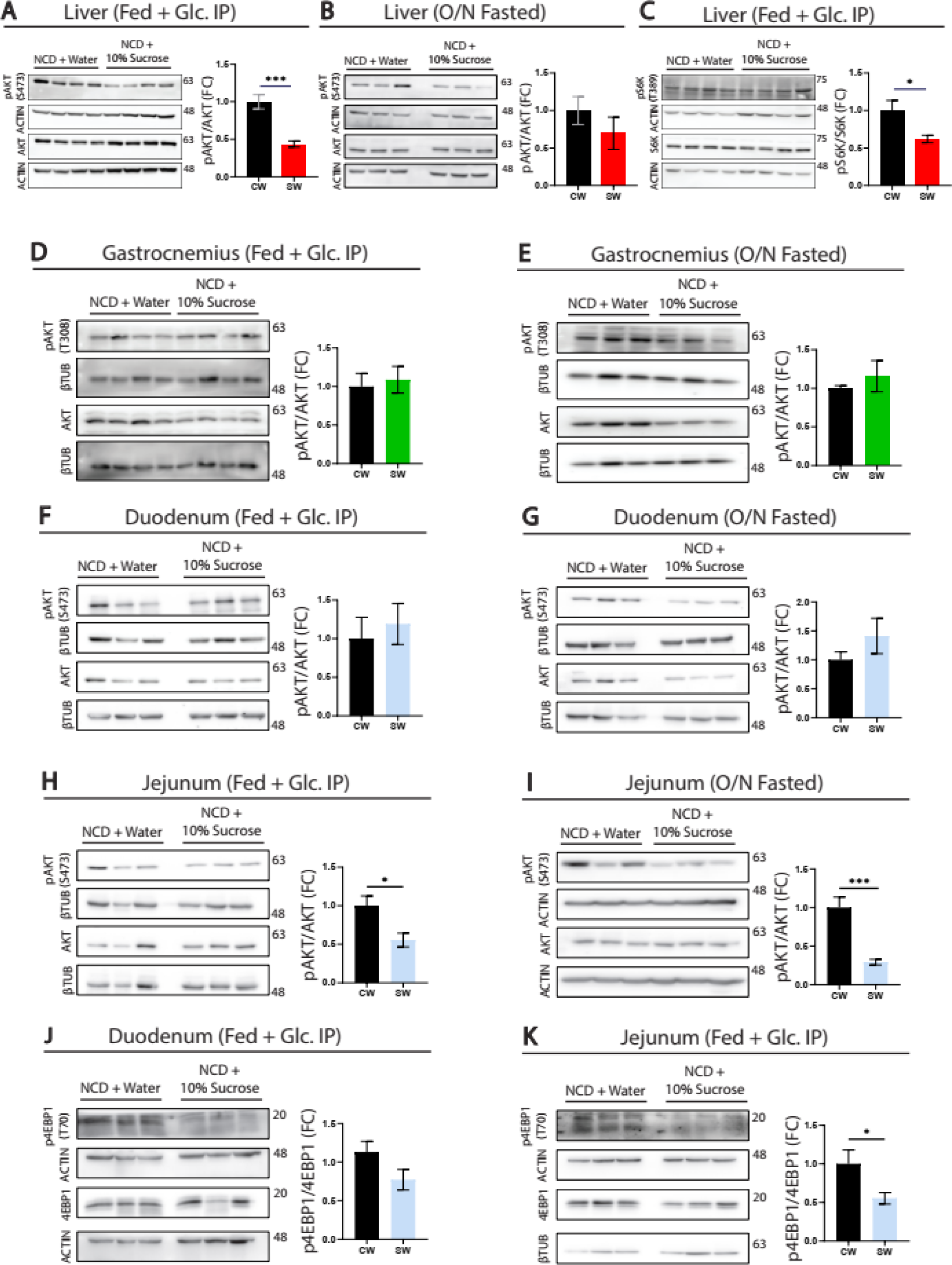
Differential regulation of nutrient and metabolic sensing/signaling upon sucrose overfeeding. (A-C) Representative immunoblots from the lysate of livers for (A-B) pS473-AKT and (C) pS6K(T389) from CW and SW mice, as indicated. (D,E) Representative immunoblot from the lysate of gastrocnemius for pT308-AKT from (D) ad-libitum fed and (E) 12hr/overnight fasted CW and SW mice. (F,G) Representative immunoblots from the lysate of duodenum for p-S473 AKT in (F) ad-libitum fed and (G) 12hr/overnight fasted CW and SW mice. (H,I) Representative immunoblots from the lysate of jejunum for p-S473 AKT in (H) ad-libitum fed and (I) 12hr/overnight fasted CW and SW mice. (J,K) Representative immunoblots for pT70-4EBP1 from lysates of (J) Duodenum and (K) Jejunum of ad-libitum fed CW and SW mice. CW - Control mice fed with NCD + Water, SW - Sucrose overfed mice fed with NCD + 10% Sucrose soln. Intraperitoneal glucose was administered to the ad-libitum fed group to assess insulin responsive molecular changes in a ‘postprandial state’. Experimental and technical repeats; N=2, n=3. Data represented as Mean ± SEM and analysed by t-test. P-value of 0.05 was considered significant. *P ≤ 0.05; **P ≤ 0.01; ***P ≤ 0.001.

Emerging evidence in the field has brought to the fore the dissimilarities with regards to insulin resistance in the muscle and liver that contributes to the onset and progression of diabetes. While in a late diabetic state reduced insulin signaling in both these tissues drives the pathology of the disease, it has also become apparent that during early stages this is not the case, depending on the context. Given this, it was interesting to find unaltered pAkt-T308 levels in the gastrocnemius of mice that consumed sucrose water irrespective of whether they were in the ad-lib fed state (Figure 5D) or an overnight starved state (Figure 5E). This hinted at a tissue specific rewiring of Insulin signaling in response to 10% sucrose consumption wherein insulin resistance in the liver possibly precedes that in the skeletal muscle [54,70,71].

In contrast to other tissues, nutrient signaling is less characterized in the small intestine particularly in a region-specific manner, even though it has been speculated to be bi-directionally associated with nutrient absorption. It was interesting to note that while sucrose consumption did not seem to affect pS473-AKT in the duodenum (Figures 5F, 5G), there was a robust decrease in the jejunum (Figures 5H, 5I) in sucrose fed mice. Notably, this reduction was independent of whether the mice were starved for 12-hours or were administered with intraperitoneal glucose, hinting at region-specific insulin signaling in the intestine (discussed later). TOR signaling was reduced in both duodenum and jejunum of sucrose overfed mice in the ad-libitum fed state, as indicated by reduced levels of T70-phosphorylation of 4EBP1 (Figure 5J,5K).

Together these results illustrate tissue and fed/fasted dependent changes in metabolic sensing which were hitherto unknown and highlights the differential responses elicited by sucrose consumption across organ-systems.

### Sucrose overfeeding leads to altered muscle mitochondrial functions

Dietary inputs have been shown to remodel mitochondrial functions, which together contribute to impaired metabolic homeostasis at an organismal level [19,41,42,44,47,72,73]. Others and we have earlier demonstrated that dysfunctional mitochondria drive the onset and progression of insulin resistance and dyslipidemia [46,74–76]. Therefore, we hypothesized that a chronic sucrose overfeeding paradigm would also result in changes in mitochondrial activity. To test this, we isolated mitochondria from liver and muscle, following 12-hour starvation from control and sucrose fed mice, and analyzed their functions systematically. Notably, besides baseline oxygen consumption rates, we also assessed state-2, state-3, state-3u and state-4 respirations by providing specific substrates, as indicated. In addition to providing a deconvoluted diagnostic assessment, this approach also indicates metabolic flexibility and potential rewiring of mitochondria to utilize different nutrient sources for energy homeostasis.

The mitochondria isolated from the livers of sucrose fed mice showed higher oxygen consumption rates across states -2, -3 and -3u only in the presence of palmitoyl-carnitine + malate substrate (Figures 6C, 6D). No such change was observed in the presence of Pyruvate + Malate and Succinate + Rotenone (Figures 6A, 6B, 6D) (Figure 6A-D). However, it was interesting to find that these mitochondria produced more ROS (Figure S6G) when fueled with all the key substrates and also seemed to impinge on ATP coupled respiration, as shown in Figures S6C-D.

**Figure 6:**
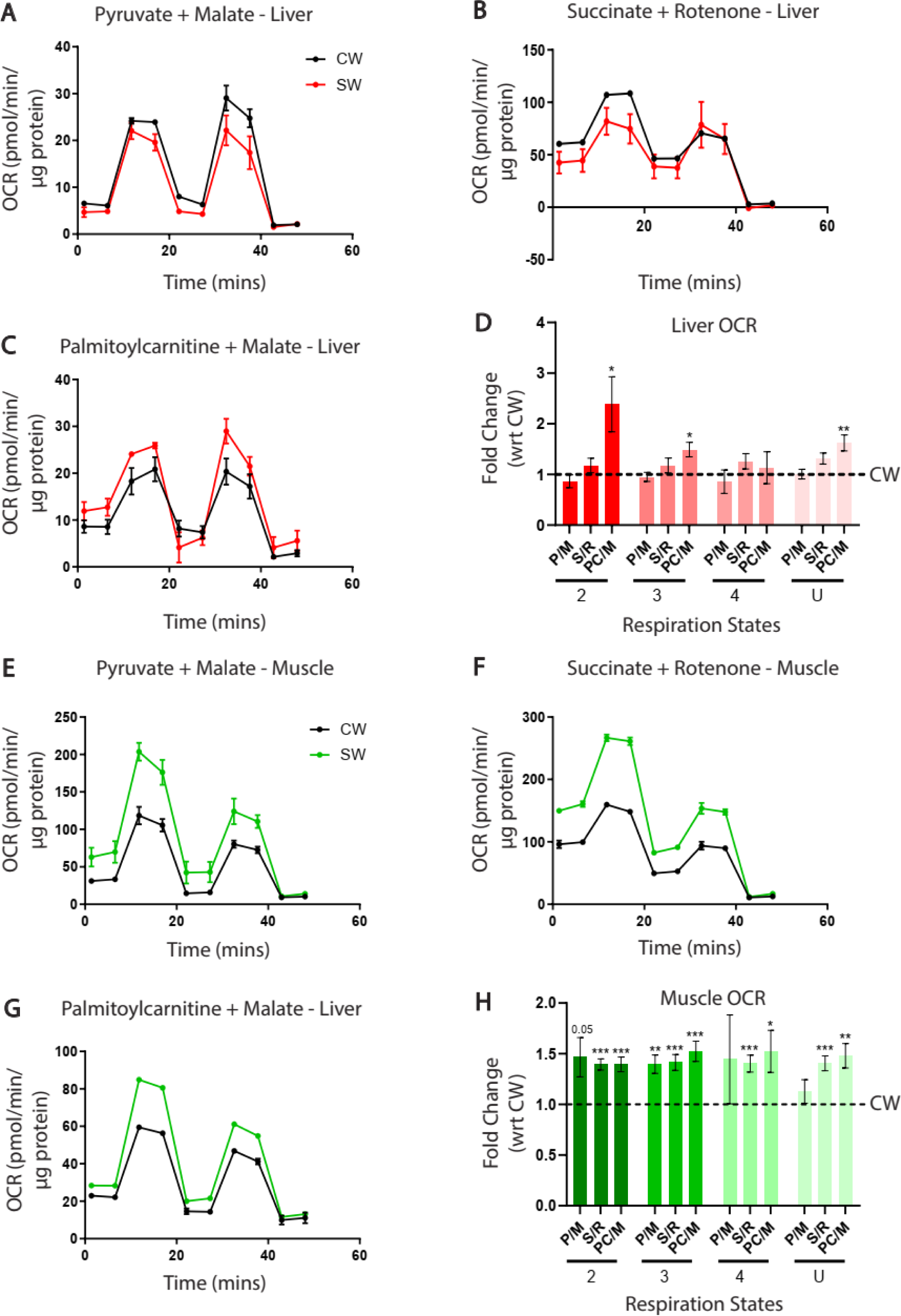
Energetic Status of Liver and Muscle Mitochondria. (A-C) Oxygen Consumption Rate (OCR) of mitochondria isolated from liver in the presence of (A) Pyruvate + Malate (P/M), (B) Succinate + Rotenone (S/R) and (C) Palmitoyl Carnitine + Malate (PC/M). (D) State-2, -3, -4 and -U respiration of mitochondria isolated from livers of SW mice in presence of P/M, S/R, PC/M and expressed as fold change with respect to liver mitochondrial OCR of CW mice. (E-G) Oxygen Consumption Rate (OCR) of mitochondria isolated from skeletal muscle (quadriceps) in the presence of (E) Pyruvate + Malate (P/M) (F) Succinate + Rotenone (S/R) and (G) Palmitoyl Carnitine + Malate (PC/M). (H) State-2, -3, -4 and -U respiration of mitochondria isolated from skeletal muscle (quadriceps) of SW mice in presence of P/M, S/R, PC/M and expressed as fold change with respect to muscle mitochondrial OCR of CW mice. Equal amounts of intact mitochondria were used for these assays as described in the Methods section. Experimental and technical repeats; N=2-3, n=3. CW - Control mice fed with NCD + Water, SW - Sucrose overfed mice fed with NCD + 10% Sucrose soln. Data represent Mean ± SEM; p<0.05(*), p<0.01(**), p<0.001(***). Statistical significance was calculated using t-test.

In contrast, mitochondria isolated from the muscle of sucrose-fed mice showed stark changes with a significant increase in oxygen consumption rates across state-2, -3, -3u and -4 for Succinate & Palmitoyl-Carnitine + Malate substrates (Figures 6F, 6G). However, Pyruvate + Malate respiration showed a significant increase only in state 3 and trends toward increase in other states (Figure 6E). Unlike in the liver, we did not observe any effect on ROS production from mitochondria (Figure S6H) isolated from muscles with no significant change in ATP coupled respiration (Figures S6E-F). Together, these results clearly suggest that there is a tissue-specific alteration of mitochondrial activity in response to chronic sucrose overfeeding.

### Intestinal nutrient transporter expression is altered in sucrose fed mice

Results described above point towards metabolic sensing and/or metabolite driven emergence of pathophysiology in response to 10% sucrose water consumption. This motivated us to investigate if molecular factors that govern nutrient absorption and transport were affected. This analysis was important to establish a potential role of intestinal macronutrient transporters as causal factors underlying the changes associated with diet-mediated physiological dys-homeostasis. Therefore, we exhaustively profiled the expression of glucose, amino-acid and fat transporters in the duodenum, jejunum and ileum along with enzymes/factors that impinge on nutrient absorption.

Sucrose is a disaccharide, which cannot be transported across the intestinal villi unless acted upon by glycosidases that generate glucose and fructose. Sucrase isomaltase is one of the predominant glycosidases in the intestine and impacts bulk absorption of monosaccharides obtained from sucrose. It was interesting to find that its transcript levels were significantly enhanced following sucrose consumption in the duodenum and jejunum (Figures 7A, 7E) although the latter is supposed to have lower isomaltase expression, since bulk of glucose absorption happens at the proximal region.

**Figure 7:**
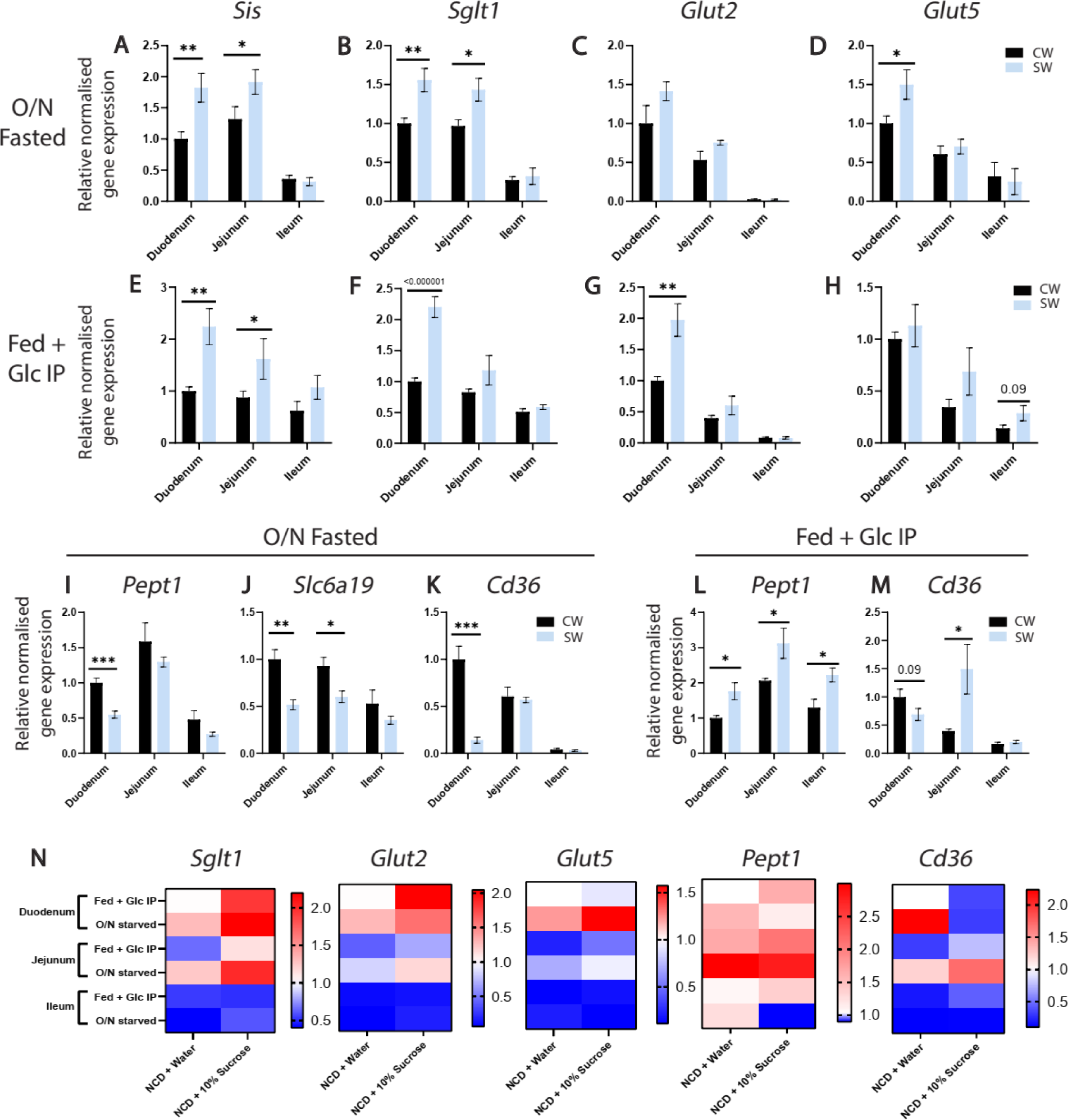
Intestinal nutrient transporter expression is altered in sucrose fed mice. (A-D) Relative intestinal (Duodenum, Jejunum and Ileum) gene expression analysis of CW and SW mice under 12hr/overnight fasted conditions. (A) Sis (Sucrose hydrolysis), (B) Sglt1 (glucose uptake from lumen), (C) Glut2 (glucose and fructose export from enterocytes), (D) Glut5 (fructose uptake). (E-H) Relative intestinal (Duodenum, Jejunum and Ileum) gene expression analysis of CW and SW mice under fed conditions. (E) Sis, (F) Sglt1, (G) Glut2, (H) Glut5. (I-K) Relative intestinal (Duodenum, Jejunum and Ileum) gene expression analysis of CW and SW mice under 12hr/overnight fasted conditions. (I) Pept1 (oligopeptide uptake), (J) Slc6a19 (branched-chain amino acid uptake), (K) Cd36 (long-chain fatty acid uptake). (L,M) Relative intestinal (Duodenum, Jejunum and Ileum) gene expression analysis of CW and SW mice under fed conditions. (L) Pept1 and (M) Cd36. Data represented as fold change with respect to expression in CW Duodenum. (N) Pair-wise intestinal (Duodenum, Jejunum and Ileum) gene expression analysis for macronutrient transporters CW and SW mice in fed and 12hr/overnight starved states. Data represented as fold change with respect to expression in Duodenum of ad-lib fed CW mice. CW - Control mice fed with NCD + Water, SW - Sucrose overfed mice fed with NCD + 10% Sucrose soln. Intraperitoneal glucose was administered to the ad-libitum fed group to assess insulin responsive molecular changes in a ‘postprandial state’. Experimental and technical repeats; N=2, n=3-4. Data represented as Mean ± SEM and analysed by (A-M) t-test and (N) two-way ANOVA. P-value of 0.05 was considered significant. *P ≤ 0.05; **P ≤ 0.01; ***P ≤ 0.001.

Albeit an earlier study has reported elevated expression of the major apical glucose transporter Sglt1 following 6% sucrose overfeeding [39], if its levels are altered across the intestinal regions and under fed/fasted states has not been addressed. We found increased expression of Sglt1 in the duodenum and jejunum, irrespective of the physiological state at which the mice were euthanized (Figures 7B, 7F). However, levels of Sglt1 transcripts in the ileum were comparable between control and sucrose cohorts.

Following apical absorption of glucose, its transport across the intestinal barrier is dependent upon glucose-6-phosphatase (G6Pc) and Glut2, a glucose transporter predominantly present on the baso-lateral surface. Given this, we found nearly overlapping changes in the expression of G6Pc (Figures 3N, 4H) and Glut2 (Figures 7C, 7G) mRNAs in the duodenum, jejunum and ileum. Similar to the Sglt1 expression pattern, Glut2 was robustly induced in the duodenum with a trend towards increase in the ileum in sucrose cohorts. The duodenum of sucrose overfed mice also showed an increase in the expression of the apical fructose transporter Glut5 following overnight starvation (Figure 7D). There was a small trend of increase in duodenal Glut5 expression observed in the fed state as well (Figure 7H). Correlative and robust increase in the expression of Sis, Sglt1, Glut5, G6pc and Glut2 posited ‘molecular addiction’ tuned towards sustained/enhanced glucose absorption/transport across the intestine.

We next wanted to investigate whether sucrose consumption also impacted the expression of other macronutrient transporters viz for amino-acid/oligopeptides and fats. Pept1 is one of the well-studied oligopeptide transporters, which contributes to bulk uptake of dietary proteins. Its expression in the Duodenum of sucrose-overfed mice was significantly reduced following 12-hour starvation (Figure 7I). Indicating a general reduction of amino-acid transporter levels as a consequence of sucrose overfeeding, transcripts of Slc6a19 and Slc1a5, which together transport branched chain and neutral amino acids, were significantly downregulated in a region-specific manner in response to 12-hour starvation (Figures 7J and S7B). However, we also noted an increase in the expression of Pept1 across all three regions in the fed state (Figure 7L).

CD36 is a ubiquitously expressed membrane protein involved in long-chain fatty acid transport. CD36 mRNA was dramatically reduced in the duodenum following 12-hour starvation (Figure 7K) and showed a trend towards decrease in ad-libitum-fed conditions (Figure 7M). Similarly, transcript abundance of Fabp2 (Figure S7A), the other key fatty acid transporter, was consistently downregulated in the duodenum of sucrose-fed cohorts.

In order to specifically unveil the differences between fed and starved states, if any, in the transcript levels of macronutrient transporters, we carried out pair-wise comparison across all the three regions of the small intestine. Besides finding sucrose consumption dependent reprogramming, expression of glucose and amino acid transporters were higher in the starved state in the proximal intestine (Figure 7N). This pattern was akin to anticipatory expression possibly to facilitate bulk macronutrient uptake upon feeding and consistent with emerging findings in the field [77–79].

Corroborating heightened Sis expression, we found elevated sucrose hydrolysis in the duodenal lysates of ad-lib fed mice (Figure 8A), which provided functional evidence to support our hypothesis of increased carbohydrate uptake following chronic consumption of sucrose water. This was further substantiated by an increase in apical SGLT1 protein levels in the duodenum of sucrose overfed mice under ad-libitum fed conditions (Figure 8B). Interestingly, this rewiring was restricted to the duodenum, with no effects in the jejunum (Figures 8C,8D). Staining with Giemsa, which is considered as a read out for mucopolysaccharides and hence of goblet cells, revealed differences between control and sucrose fed mice in the jejunum (Figures 8F, S7D, S7E) and ileum (Figures S7D, S7E, S7F). Collectively, these illustrated differential effects of 10% sucrose water consumption on the intestinal epithelium at molecular, cellular and functional levels in a region-specific manner.

**Figure 8:**
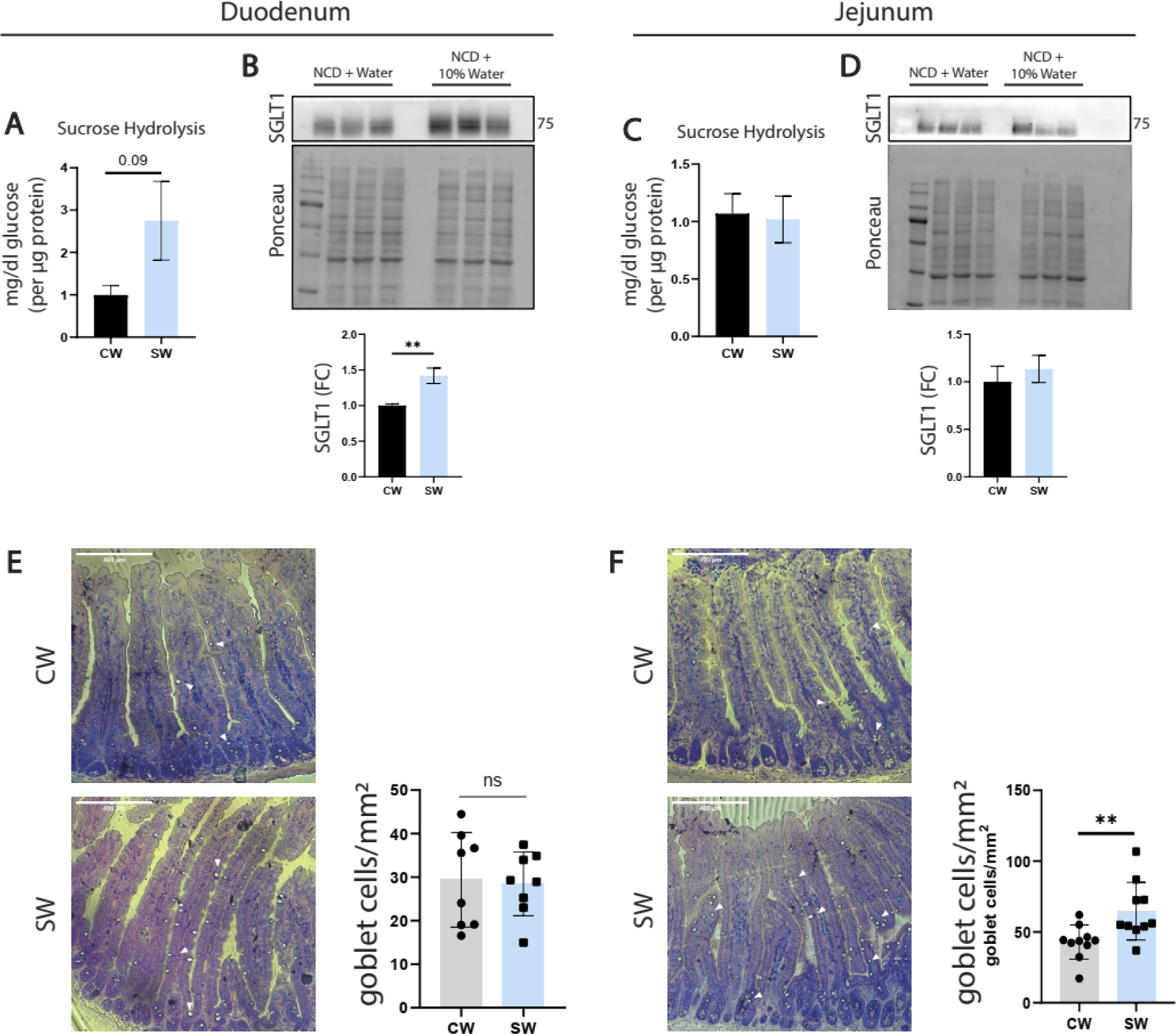
Ad-libitum fed mice show elevated duodenal hexose uptake upon chronic sucrose overfeeding. (A) Sucrose hydrolysis in duodenum of CW and SW mice under fed conditions. (B) Representative immunoblot for SGLT1 transporter levels in the apical membrane fraction of Duodenum from CW and SW mice. (C) Sucrose hydrolysis in jejunum of CW and SW mice under fed conditions. (D) Representative immunoblot for SGLT1 transporter levels in the apical membrane fraction of Jejunum from CW and SW mice. (E,F) Representative histological sections and quantification of goblet cell population in (E) Duodenum and (F) Jejunum of sucrose overfed mice. Sections obtained from fed CW and SW mice and stained with Giemsa for visualization of goblet cells. CW - Control mice fed with NCD + Water, SW - Sucrose overfed mice fed with NCD + 10% Sucrose soln. Experimental and technical repeats; N=2, n=3 for sucrose hydrolysis and SGLT1 level quantification. Data represented as Mean ± SEM and analysed by t-test (A-D). Intraperitoneal glucose was administered to the ad-libitum fed group to assess insulin responsive molecular changes in a ‘postprandial state’. Experimental and technical repeats N=2, n= 8-10 for goblet cell data, represented as Mean ± SD and analysed by t-test (E,F). P-value of 0.05 was considered significant. *P ≤ 0.05; **P ≤ 0.01; ***P ≤ 0.001.

## Discussion

Sucrose over consumption, globally, has been proposed as a leading cause of metabolic dysfunction. Despite this, very few reports have provided molecular, histological and physiological insights across organ-systems, especially in response to sucrose water intake, which reasonably mimics human consumption. Given that commonly consumed sugar-sweetened beverages typically contain 10-15% sucrose, we have exhaustively investigated underlying mechanisms, which are causally or consequently affected, and collectively contribute towards physiological mal-effects of 10% sucrose water feeding, at a systems level. Notably, we demonstrate how these effects are mediated through non-overlapping and distinct pathways that operate across the organ systems.

Most of our current understanding of mechanisms that cause disease-like states following chronic high calorie diets, including sucrose consumption, are limited to changes observed in an ad-libitum fed state. This has led to biased interpretations and incomplete assessment of physiological deficits. Anabolic and catabolic balance is well known to be crucial for homeostasis and emerging reports including from our group have highlighted the importance of elucidating mechanisms in both fed and fasted states. In this context, our study provides a comprehensive picture of pathways/factors that are affected in a physiological state-specific manner following chronic 10% sucrose water consumption (Figure 9).

**Figure 9:**
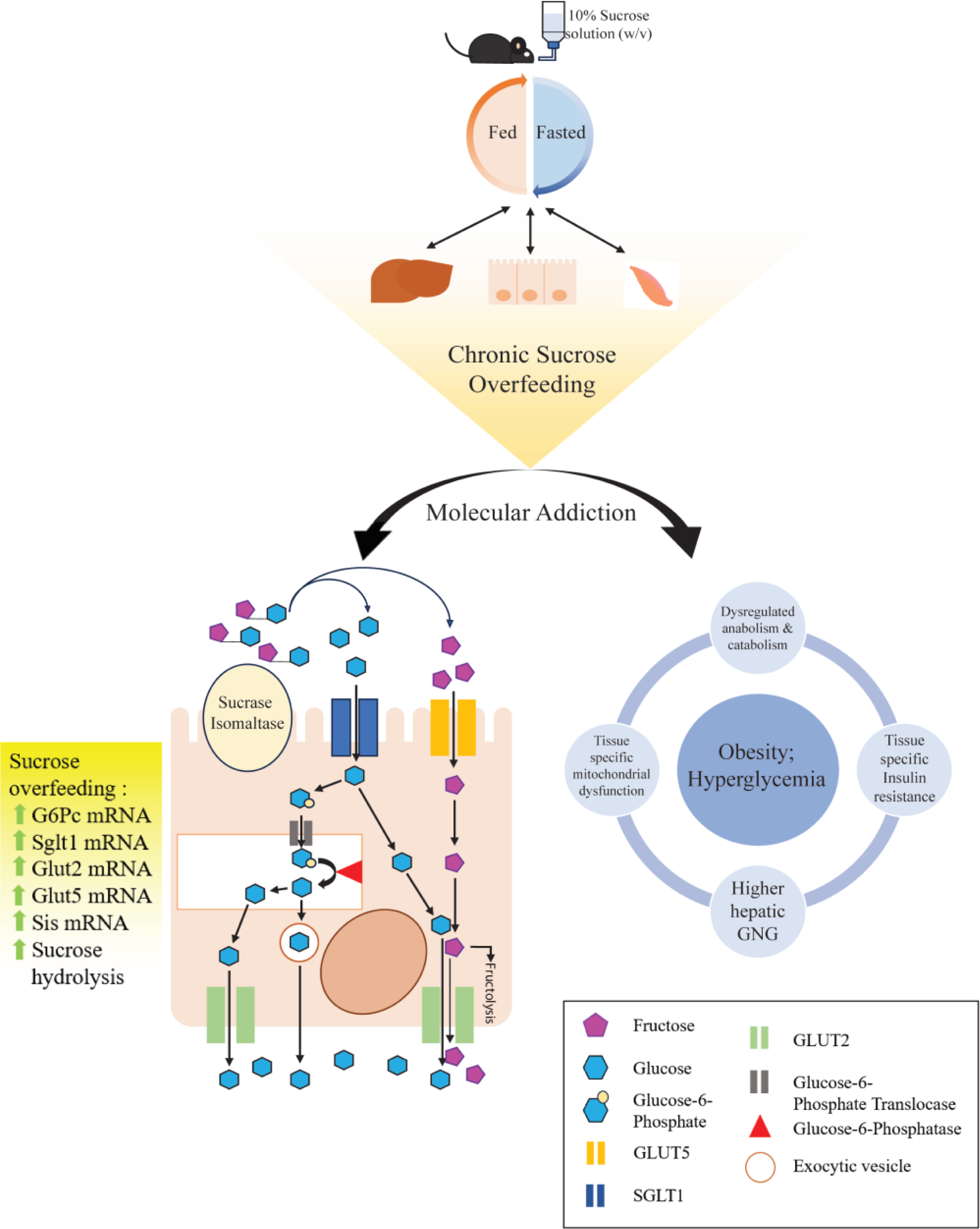
Model for ‘molecular addiction’ to sucrose. Schematic representation of ‘molecular addiction’ of the small intestine to sucrose. The increased sucrose hydrolysis in the proximal small intestine leads to increased uptake of glucose and fructose into the enterocytes through SGLT1 and GLUT5 respectively. Glucose and fructose can then get exported from the basolateral surface through GLUT2. In the fed state, glucose and fructose can enhance the DNA binding capability of CHREBP which can drive transcriptional upregulation of Glucose-6-Phosphatase (G6Pc). The G6Pc can also drive transepithelial transport of glucose wherein cytosolic Glucose-6-phosphate is transported to the ER lumen and acted upon by G6Pc to release free glucose. This glucose can then be exported from the basolateral surface either through GLUT2 or by means of exocytosis. Sucrose overfeeding thus employs both phophorylation-dependent and -independent modes of transepithelial glucose transport in the small intestine leading to higher levels of glucose reaching the circulation.

In consonance with the literature [31,39,45,49,70], and as anticipated, sucrose water consumption led to weight gain and impaired glucose homeostasis. Although not surprising, our findings indicate that both insulin resistance and hepatic gluconeogenesis contribute to elevated blood glucose levels in sucrose-water fed mice. While reports employing very high sucrose consumption have shown drastic changes in hepatic gene expression [29], our results clearly illustrate that 10% sucrose water impacts hepatic gene expression in a state specific manner. While sucrose overfeeding does not impinge on the expression of key metabolic genes in the liver of overnight fasted mice, genes regulating lipid homeostasis do show perturbed expression in the livers of sucrose fed mice in the ad-lib fed state. Nonetheless, significant reduction in insulin signaling and enhanced mitochondrial ROS production posits hepatic insufficiency in glucose metabolism is independent of gene-expression changes. This is important since a majority of efforts to tackle metabolic diseases are aimed at targeting transcription in the liver.

Muscles have been regarded as the major ‘consumers’ of glucose and are invariably associated with excess carbohydrate mediated onset of diabetes and obesity. Others and we have shown evolutionarily conserved mechanisms that impinge on the mitochondria as being causal to metabolic deficits in the muscles, including during aging [74,80,81]. In our efforts to tease out multi-organ involvement, we have found significant alterations in mitochondrial functions and biased substrate oxidation in sucrose fed mice.

Several studies have reported dysbiosis of the intestine, especially of the colon, in response to high calorie intake [18,20,82–86]. However, even though the small intestine is one of the largest organs and is pivotal to nutrient absorption, very little is known about its involvement in chronic dietary paradigms. Specifically, duodenal, jejunal and ileal mechanisms that are differentially affected following 10% sucrose water consumption, and in a fed-fast dependent manner, have not been well characterized. In this regard, our findings reveal region-specific differential impact of sucrose feeding, which were hitherto unknown. It is important to note that while expression of glucose transporters like Sglt1, Glut5 and Glut2 were elevated, we found dampened expression of amino-acid and fatty-acid transporters (Pept1, Cd36, Slc1a5, Slc6a19, Fabp2). This was associated with increased villi length, and correlated with enhanced proliferation. Changes in villi architecture were limited to the proximal intestine where carbohydrate is maximally absorbed and in the absence of any other gross abnormalities, indicate an active/direct involvement of epithelial cells. Taken together, these clearly hinted at rewiring of intestinal epithelium to sustain carbohydrate absorption, which could also affect the uptake of other macronutrients.

Besides transporters, uptake of glucose and fructose in the proximal intestine is dependent upon enzymes that breakdown sucrose in the lumen and those within enterocytes that enable basolateral transport. Consistent with this, we did indeed find robust changes in the expression of sucrase isomaltase, aldolase and triokinase in response to 10% sucrose water consumption. This was further substantiated by elevated G6Pase levels and reduced insulin signaling, which are essential for ‘delivering’ glucose at the basolateral surface, possibly by reducing intestinal utilization of glucose. These findings posit a scenario wherein sucrose water consumption leads to intestinal addiction to sucrose, which could be the primary driver of the pathophysiology.

Excess carbohydrate intake has long been known to cause physiological impairment and a public health crisis, which has increased owing to over consumption of sugar-sweetened beverages. Our findings provide a comprehensive picture of the molecular and cellular mechanisms that are associated with metabolic deficits following chronic consumption of human relevant sucrose water. In addition, these invoke a multi-organ cross-talk, which is likely driven by intestinal mal-absorption of glucose and fructose. Besides bringing to the fore the need to further our understanding of mechanisms that rewire nutrient uptake and utilization across organ-systems, this study will aid in developing strategies to counter this ‘molecular addiction’.

### CRediT authorship contribution statement

**Saptarnab Ganguly :** Investigation, Data Curation, Formal Analysis, Methodology, Validation, Writing - original draft, **Tandrika Chattopadhyay :** Investigation, Data Curation, Formal Analysis, Methodology, Writing - review & editing, **Rubina Kazi :** Project Administration, **Souparno Das :** Investigation, Data Curation, Formal Analysis, Methodology, Validation, Project Administration, **Bhavisha Malik :** Investigation, Data Curation, Formal Analysis, Methodology, Validation, **Uthpala M.L. :** Investigation, Data Curation, Formal Analysis, Validation, **Padmapriya S. Iyer :** Investigation, Data Curation, Formal Analysis, Validation, **Mohit Kashiv :** Investigation, Methodology, **Anshit Singh :** Investigation, **Amita Ghadge :** Investigation, **Shyam D. Nair :** Project Administration, **Mahendra S. Sonawane :** Conceptualization, Funding Acquisition, Resources, Supervision, Writing - review & editing, **Ullas Kolthur-Seetharam :** Conceptualization, Funding Acquisition, Resources, Supervision, Writing - original draft, Writing - review & editing.

## Acknowledgements

We are grateful to IISER Pune Animal House for providing us C57BL/6N mice. We particularly thank Dr. Kalidas N. Kohale, Dr. Shital Suryavanshi, Dr. Gopalakrishna R, Chetan Sable, Kanakam Kishore, TIFR Mumbai, TIFR Hyderabad and IISER Pune animal house staff for their invaluable help with animal experiments. We extend our gratitude to the members of UK lab and MS lab for their critical inputs and discussions during the study.

## Conflict of Interest

The authors declare that they have no conflict of interest

## Funding

This research has been supported by the following funding sources : TIFR/DAE (19P0116), TIFR/DAE(19P0911), TIFR/DAE(19P2143) and Department of Biotechnology (BT/PR29878/PFN/20/1431/2018).

## Notes

### Competing Interest Statement

Advanced Research Unit on Metabolism Development and Aging (ARUMDA) in TIFR has recieved CSR funding from Hindustan Unilever Ltd. This funding/interaction has not conflict of interest for the findings described in this manuscript.

## Bibliography

[1] Schmitz J, Preiser H, Maestracci D, Ghosh BK, Cerda JJ, Crane RK. Purification of the human intestinal brush border membrane. Biochimica et Biophysica Acta (BBA) - Biomembranes 1973;323:98–112. 10.1016/0005-2736(73)90434-3.

[2] Lara-Castor L, Micha R, Cudhea F, Miller V, Shi P, Zhang J, et al. Sugar-sweetened beverage intakes among adults between 1990 and 2018 in 185 countries. Nat Commun 2023;14:5957. 10.1038/s41467-023-41269-8.

[3] Singh GM, Micha R, Khatibzadeh S, Lim S, Ezzati M, Mozaffarian D, et al. Estimated Global, Regional, and National Disease Burdens Related to Sugar-Sweetened Beverage Consumption in 2010. Circulation 2015;132:639–66. 10.1161/CIRCULATIONAHA.114.010636.

[4] Della Corte K, Fife J, Gardner A, Murphy BL, Kleis L, Della Corte D, et al. World trends in sugar-sweetened beverage and dietary sugar intakes in children and adolescents: a systematic review. Nutr Rev 2021;79:274–88. 10.1093/nutrit/nuaa070.

[5] Malik VS, Hu FB. The role of sugar-sweetened beverages in the global epidemics of obesity and chronic diseases. Nat Rev Endocrinol 2022;18:205–18. 10.1038/s41574-021-00627-6.

[6] Rodríguez LA, Madsen KA, Cotterman C, Lustig RH. Added sugar intake and metabolic syndrome in US adolescents: cross-sectional analysis of the National Health and Nutrition Examination Survey 2005-2012. Public Health Nutr 2016;19:2424–34. 10.1017/S1368980016000057.

[7] Demaria TM, Crepaldi LD, Costa-Bartuli E, Branco JR, Zancan P, Sola-Penna M. Once a week consumption of Western diet over twelve weeks promotes sustained insulin resistance and non-alcoholic fat liver disease in C57BL/6 J mice. Sci Rep 2023;13:3058. 10.1038/s41598-023-30254-2.

[8] Distefano JK, Gerhard GS. Effects of dietary sugar restriction on hepatic fat in youth with obesity. Minerva Pediatr (Torino) 2023. 10.23736/S2724-5276.23.07209-9.

[9] Nash MJ, Dobrinskikh E, Janssen RC, Lovell MA, Schady DA, Levek C, et al. Maternal Western diet is associated with distinct preclinical pediatric NAFLD phenotypes in juvenile nonhuman primate offspring. Hepatol Commun 2023;7:e0014. 10.1097/HC9.0000000000000014.

[10] Renault KM, Carlsen EM, Nørgaard K, Nilas L, Pryds O, Secher NJ, et al. Intake of sweets, snacks and soft drinks predicts weight gain in obese pregnant women: detailed analysis of the results of a randomised controlled trial. PLoS ONE 2015;10:e0133041. 10.1371/journal.pone.0133041.

[11] Maslova E, Halldorsson TI, Astrup A, Olsen SF. Dietary protein-to-carbohydrate ratio and added sugar as determinants of excessive gestational weight gain: a prospective cohort study. BMJ Open 2015;5:e005839. 10.1136/bmjopen-2014-005839.

[12] Weijs PJ m, Kool LM, van Baar NM, van der Zee SC. High beverage sugar as well as high animal protein intake at infancy may increase overweight risk at 8 years: a prospective longitudinal pilot study. Nutr J 2011;10:95. 10.1186/1475-2891-10-95.

[13] Reiser S, Bohn E, Hallfrisch J, Michaelis OE, Keeney M, Prather ES. Serum insulin and glucose in hyperinsulinemic subjects fed three different levels of sucrose. Am J Clin Nutr 1981;34:2348–58. 10.1093/ajcn/34.11.2348.

[14] Commerford SR, Ferniza JB, Bizeau ME, Thresher JS, Willis WT, Pagliassotti MJ. Diets enriched in sucrose or fat increase gluconeogenesis and G-6-Pase but not basal glucose production in rats. Am J Physiol Endocrinol Metab 2002;283:E545–55. 10.1152/ajpendo.00120.2002.

[15] Hulman S, Falkner B. The effect of excess dietary sucrose on growth, blood pressure, and metabolism in developing Sprague-Dawley rats. Pediatr Res 1994;36:95–101. 10.1203/00006450-199407001-00017.

[16] Hasegawa Y, Chen S-Y, Sheng L, Jena PK, Kalanetra KM, Mills DA, et al. Long-term effects of western diet consumption in male and female mice. Sci Rep 2020;10:14686. 10.1038/s41598-020-71592-9.

[17] De Groef S, Wilms T, Balmand S, Calevro F, Callaerts P. Sexual Dimorphism in Metabolic Responses to Western Diet in Drosophila melanogaster. Biomolecules 2021;12. 10.3390/biom12010033.

[18] Romualdo GR, Valente LC, Sprocatti AC, Bacil GP, de Souza IP, Rodrigues J, et al. Western diet-induced mouse model of non-alcoholic fatty liver disease associated with metabolic outcomes: Features of gut microbiome-liver-adipose tissue axis. Nutrition 2022;103–104:111836. 10.1016/j.nut.2022.111836.

[19] Liu I-F, Lin T-C, Wang S-C, Yen C-H, Li C-Y, Kuo H-F, et al. Long-term administration of Western diet induced metabolic syndrome in mice and causes cardiac microvascular dysfunction, cardiomyocyte mitochondrial damage, and cardiac remodeling involving caveolae and caveolin-1 expression. Biol Direct 2023;18:9. 10.1186/s13062-023-00363-z.

[20] Martínez Leo EE, Segura Campos MR. Effect of ultra-processed diet on gut microbiota and thus its role in neurodegenerative diseases. Nutrition 2020;71:110609. 10.1016/j.nut.2019.110609.

[21] Yang M, Qi X, Li N, Kaifi JT, Chen S, Wheeler AA, et al. Western diet contributes to the pathogenesis of non-alcoholic steatohepatitis in male mice via remodeling gut microbiota and increasing production of 2-oleoylglycerol. Nat Commun 2023;14:228. 10.1038/s41467-023-35861-1.

[22] Musial B, Vaughan OR, Fernandez-Twinn DS, Voshol P, Ozanne SE, Fowden AL, et al. A Western-style obesogenic diet alters maternal metabolic physiology with consequences for fetal nutrient acquisition in mice. J Physiol (Lond) 2017;595:4875–92. 10.1113/JP273684.

[23] Hirahatake KM, Meissen JK, Fiehn O, Adams SH. Comparative effects of fructose and glucose on lipogenic gene expression and intermediary metabolism in HepG2 liver cells. PLoS ONE 2011;6:e26583. 10.1371/journal.pone.0026583.

[24] Jiang L, Wang Q, Yu Y, Zhao F, Huang P, Zeng R, et al. Leptin contributes to the adaptive responses of mice to high-fat diet intake through suppressing the lipogenic pathway. PLoS ONE 2009;4:e6884. 10.1371/journal.pone.0006884.

[25] Daly ME, Vale C, Walker M, Littlefield A, George K, Alberti M, et al. Acute fuel selection in response to high-sucrose and high-starch meals in healthy men. Am J Clin Nutr 2000;71:1516–24. 10.1093/ajcn/71.6.1516.

[26] Thresher JS, Podolin DA, Wei Y, Mazzeo RS, Pagliassotti MJ. Comparison of the effects of sucrose and fructose on insulin action and glucose tolerance. Am J Physiol Regul Integr Comp Physiol 2000;279:R1334–40. 10.1152/ajpregu.2000.279.4.R1334.

[27] Ivić V, Zjalić M, Blažetić S, Fenrich M, Labak I, Scitovski R, et al. Elderly rats fed with a high-fat high-sucrose diet developed sex-dependent metabolic syndrome regardless of long-term metformin and liraglutide treatment. Front Endocrinol (Lausanne) 2023;14:1181064. 10.3389/fendo.2023.1181064.

[28] Rasool S, Geetha T, Broderick TL, Babu JR. High fat with high sucrose diet leads to obesity and induces myodegeneration. Front Physiol 2018;9:1054. 10.3389/fphys.2018.01054.

[29] Schultz A, Barbosa-da-Silva S, Aguila MB, Mandarim-de-Lacerda CA. Differences and similarities in hepatic lipogenesis, gluconeogenesis and oxidative imbalance in mice fed diets rich in fructose or sucrose. Food Funct 2015;6:1684–91. 10.1039/c5fo00251f.

[30] Fuente-Martín E, Granado M, García-Cáceres C, Sanchez-Garrido MA, Frago LM, Tena-Sempere M, et al. Early nutritional changes induce sexually dimorphic long-term effects on body weight gain and the response to sucrose intake in adult rats. Metab Clin Exp 2012;61:812–22. 10.1016/j.metabol.2011.11.003.

[31] Stephenson EJ, Stayton AS, Sethuraman A, Rao PK, Meyer A, Gomes CK, et al. Chronic intake of high dietary sucrose induces sexually dimorphic metabolic adaptations in mouse liver and adipose tissue. Nat Commun 2022;13:6062. 10.1038/s41467-022-33840-6.

[32] Ecker A, Gonzaga TKS do N, Seeger RL, Santos MMD, Loreto JS, Boligon AA, et al. High-sucrose diet induces diabetic-like phenotypes and oxidative stress in Drosophila melanogaster: Protective role of Syzygium cumini and Bauhinia forficata. Biomed Pharmacother 2017;89:605–16. 10.1016/j.biopha.2017.02.076.

[33] Gatineau E, Savary-Auzeloux I, Migné C, Polakof S, Dardevet D, Mosoni L. Chronic Intake of Sucrose Accelerates Sarcopenia in Older Male Rats through Alterations in Insulin Sensitivity and Muscle Protein Synthesis. J Nutr 2015;145:923–30. 10.3945/jn.114.205583.

[34] de Wit NJ, Bosch-Vermeulen H, de Groot PJ, Hooiveld GJ, Bromhaar MMG, Jansen J, et al. The role of the small intestine in the development of dietary fat-induced obesity and insulin resistance in C57BL/6J mice. BMC Med Genomics 2008;1:14. 10.1186/1755-8794-1-14.

[35] Bein A, Fadel CW, Swenor B, Cao W, Powers RK, Camacho DM, et al. Nutritional deficiency in an intestine-on-a-chip recapitulates injury hallmarks associated with environmental enteric dysfunction. Nat Biomed Eng 2022;6:1236–47. 10.1038/s41551-022-00899-x.

[36] Pinho RM, Garas LC, Huang BC, Weimer BC, Maga EA. Malnourishment affects gene expression along the length of the small intestine. Front Nutr 2022;9:894640. 10.3389/fnut.2022.894640.

[37] Garas LC, Hamilton MK, Dawson MW, Wang J-L, Murray JD, Raybould HE, et al. Lysozyme-rich milk mitigates effects of malnutrition in a pig model of malnutrition and infection. Br J Nutr 2018;120:1131–48. 10.1017/S0007114518002507.

[38] Segú H, Jalševac F, Pinent M, Ardévol A, Terra X, Blay MT. Intestinal Morphometric Changes Induced by a Western-Style Diet in Wistar Rats and GSPE Counter-Regulatory Effect. Nutrients 2022;14. 10.3390/nu14132608.

[39] Wu X, Cui L, Wang H, Xu J, Zhong Z, Jia X, et al. Impact of dietary sucralose and sucrose-sweetened water intake on lipid and glucose metabolism in male mice. Eur J Nutr 2023;62:199–211. 10.1007/s00394-022-02980-2.

[40] Kim JY, Nolte LA, Hansen PA, Han DH, Kawanaka K, Holloszy JO. Insulin resistance of muscle glucose transport in male and female rats fed a high-sucrose diet. Am J Physiol 1999;276:R665–72. 10.1152/ajpregu.1999.276.3.R665.

[41] Aimaretti E, Chimienti G, Rubeo C, Di Lorenzo R, Trisolini L, Dal Bello F, et al. Different Effects of High-Fat/High-Sucrose and High-Fructose Diets on Advanced Glycation End-Product Accumulation and on Mitochondrial Involvement in Heart and Skeletal Muscle in Mice. Nutrients 2023;15. 10.3390/nu15234874.

[42] Jørgensen W, Rud KA, Mortensen OH, Frandsen L, Grunnet N, Quistorff B. Your mitochondria are what you eat: a high-fat or a high-sucrose diet eliminates metabolic flexibility in isolated mitochondria from rat skeletal muscle. Physiol Rep 2017;5. 10.14814/phy2.13207.

[43] Pagliassotti MJ, Shahrokhi KA, Moscarello M. Involvement of liver and skeletal muscle in sucrose-induced insulin resistance: dose-response studies. Am J Physiol 1994;266:R1637–44. 10.1152/ajpregu.1994.266.5.R1637.

[44] Ruiz-Ramírez A, Chávez-Salgado M, Peñeda-Flores JA, Zapata E, Masso F, El-Hafidi M. High-sucrose diet increases ROS generation, FFA accumulation, UCP2 level, and proton leak in liver mitochondria. Am J Physiol Endocrinol Metab 2011;301:E1198-207. 10.1152/ajpendo.00631.2010.

[45] Ozkan H, Yakan A. Dietary high calories from sunflower oil, sucrose and fructose sources alters lipogenic genes expression levels in liver and skeletal muscle in rats. Ann Hepatol 2019;18:715–24. 10.1016/j.aohep.2019.03.013.

[46] Cruz Hernández JH, Rosado Lomán WN, Gómez-Crisóstomo NP, De la Cruz-Hernández EN, Guzmán García LM, Gómez Gómez M, et al. High sugar but not high fat diet consumption induces hepatic metabolic disruption and up-regulation of mitochondrial fission-associated protein Drp1 in a model of moderate obesity. Arch Physiol Biochem 2023;129:233–40. 10.1080/13813455.2020.1812666.

[47] Mahler CA, Snoke DB, Cole RM, Angelotti A, Sparagna GC, Baskin KK, et al. Consuming a Linoleate-Rich Diet Increases Levels of Tetralinoleoyl Cardiolipin in Mouse Liver and Alters Hepatic Mitochondrial Respiration. J Nutr 2023. 10.1016/j.tjnut.2023.12.037.

[48] Mahmood A, Faisal MN, Khan JA, Muzaffar H, Muhammad F, Hussain J, et al. Association of a high-fat diet with I-FABP as a biomarker of intestinal barrier dysfunction driven by metabolic changes in Wistar rats. Lipids Health Dis 2023;22:68. 10.1186/s12944-023-01837-9.

[49] Kobi JBBS, Matias AM, Gasparini PVF, Torezani-Sales S, Madureira AR, da Silva DS, et al. High-fat, high-sucrose, and combined high-fat/high-sucrose diets effects in oxidative stress and inflammation in male rats under presence or absence of obesity. Physiol Rep 2023;11:e15635. 10.14814/phy2.15635.

[50] Ishimoto T, Lanaspa MA, Rivard CJ, Roncal-Jimenez CA, Orlicky DJ, Cicerchi C, et al. High-fat and high-sucrose (western) diet induces steatohepatitis that is dependent on fructokinase. Hepatology 2013;58:1632–43. 10.1002/hep.26594.

[51] Liu Y, Li M, Lv X, Bao K, Yu Tian X, He L, et al. Yes-Associated Protein Targets the Transforming Growth Factor β Pathway to Mediate High-Fat/High-Sucrose Diet-induced Arterial Stiffness. Circ Res 2022;130:851–67. 10.1161/CIRCRESAHA.121.320464.

[52] Murase T, Aoki M, Wakisaka T, Hase T, Tokimitsu I. Anti-obesity effect of dietary diacylglycerol in C57BL/6J mice: dietary diacylglycerol stimulates intestinal lipid metabolism. J Lipid Res 2002;43:1312–9. 10.1194/jlr.M200094-JLR200.

[53] Shaw E, Talwadekar M, Rashida Z, Mohan N, Acharya A, Khatri S, et al. Anabolic SIRT4 Exerts Retrograde Control over TORC1 Signaling by Glutamine Sparing in the Mitochondria. Mol Cell Biol 2020;40. 10.1128/MCB.00212-19.

[54] Chicco A, D’Alessandro ME, Karabatas L, Pastorale C, Basabe JC, Lombardo YB. Muscle lipid metabolism and insulin secretion are altered in insulin-resistant rats fed a high sucrose diet. J Nutr 2003;133:127–33. 10.1093/jn/133.1.127.

[55] Sokolović M, Wehkamp D, Sokolović A, Vermeulen J, Gilhuijs-Pederson LA, van Haaften RIM, et al. Fasting induces a biphasic adaptive metabolic response in murine small intestine. BMC Genomics 2007;8:361. 10.1186/1471-2164-8-361.

[56] Yoshikawa T, Inoue R, Matsumoto M, Yajima T, Ushida K, Iwanaga T. Comparative expression of hexose transporters (SGLT1, GLUT1, GLUT2 and GLUT5) throughout the mouse gastrointestinal tract. Histochem Cell Biol 2011;135:183–94. 10.1007/s00418-011-0779-1.

[57] Rao A, Haywood J, Craddock AL, Belinsky MG, Kruh GD, Dawson PA. The organic solute transporter alpha-beta, Ostalpha-Ostbeta, is essential for intestinal bile acid transport and homeostasis. Proc Natl Acad Sci USA 2008;105:3891–6. 10.1073/pnas.0712328105.

[58] Stine RR, Sakers AP, TeSlaa T, Kissig M, Stine ZE, Kwon CW, et al. PRDM16 Maintains Homeostasis of the Intestinal Epithelium by Controlling Region-Specific Metabolism. Cell Stem Cell 2019;25:830–845.e8. 10.1016/j.stem.2019.08.017.

[59] Deota S, Chattopadhyay T, Ramachandran D, Armstrong E, Camacho B, Maniyadath B, et al. Identification of a Tissue-Restricted Isoform of SIRT1 Defines a Regulatory Domain that Encodes Specificity. Cell Rep 2017;18:3069–77. 10.1016/j.celrep.2017.03.012.

[60] Thomas M, Bayha C, Klein K, Müller S, Weiss TS, Schwab M, et al. The truncated splice variant of peroxisome proliferator-activated receptor alpha, PPARα-tr, autonomously regulates proliferative and pro-inflammatory genes. BMC Cancer 2015;15:488. 10.1186/s12885-015-1500-x.

[61] Léveillé M, Besse-Patin A, Jouvet N, Gunes A, Sczelecki S, Jeromson S, et al. PGC-1α isoforms coordinate to balance hepatic metabolism and apoptosis in inflammatory environments. Mol Metab 2020;34:72–84. 10.1016/j.molmet.2020.01.004.

[62] Rogacka D, Piwkowska A, Audzeyenka I, Angielski S, Jankowski M. SIRT1-AMPK crosstalk is involved in high glucose-dependent impairment of insulin responsiveness in primary rat podocytes. Exp Cell Res 2016;349:328–38. 10.1016/j.yexcr.2016.11.005.

[63] Jeon S-M. Regulation and function of AMPK in physiology and diseases. Exp Mol Med 2016;48:e245. 10.1038/emm.2016.81.

[64] Yu D, Richardson NE, Green CL, Spicer AB, Murphy ME, Flores V, et al. The adverse metabolic effects of branched-chain amino acids are mediated by isoleucine and valine. Cell Metab 2021;33:905–922.e6. 10.1016/j.cmet.2021.03.025.

[65] Cantó C, Gerhart-Hines Z, Feige JN, Lagouge M, Noriega L, Milne JC, et al. AMPK regulates energy expenditure by modulating NAD+ metabolism and SIRT1 activity. Nature 2009;458:1056–60. 10.1038/nature07813.

[66] Howell JJ, Hellberg K, Turner M, Talbott G, Kolar MJ, Ross DS, et al. Metformin Inhibits Hepatic mTORC1 Signaling via Dose-Dependent Mechanisms Involving AMPK and the TSC Complex. Cell Metab 2017;25:463–71. 10.1016/j.cmet.2016.12.009.

[67] Soeters MR, Soeters PB, Schooneman MG, Houten SM, Romijn JA. Adaptive reciprocity of lipid and glucose metabolism in human short-term starvation. Am J Physiol Endocrinol Metab 2012;303:E1397–407. 10.1152/ajpendo.00397.2012.

[68] Blackmore K, Zhou W, Dailey MJ. LKB1-AMPK modulates nutrient-induced changes in the mode of division of intestinal epithelial crypt cells in mice. Exp Biol Med (Maywood) 2017;242:1490–8. 10.1177/1535370217724427.

[69] Shukla N, Kadam S, Padinhateeri R, Kolthur-Seetharam U. Continuous variable responses and signal gating form kinetic bases for pulsatile insulin signaling and emergence of resistance. Proc Natl Acad Sci USA 2021;118. 10.1073/pnas.2102560118.

[70] Chicco A, DAless M, Karabatas L, Pastorale C, Basabe J, Lombardo. Muscle Lipid Metabolism and Insulin Secretion Are Altered in Insulin-Resistant Rats Fed a High Sucrose Diet1,2 n.d.

[71] Pagliassotti MJ, Prach PA, Koppenhafer TA, Pan DA. Changes in insulin action, triglycerides, and lipid composition during sucrose feeding in rats. Am J Physiol 1996;271:R1319–26. 10.1152/ajpregu.1996.271.5.R1319.

[72] Chanseaume E, Giraudet C, Gryson C, Walrand S, Rousset P, Boirie Y, et al. Enhanced muscle mixed and mitochondrial protein synthesis rates after a high-fat or high-sucrose diet. Obesity (Silver Spring) 2007;15:853–9. 10.1038/oby.2007.582.

[73] Kutz LC, Cui X, Xie JX, Mukherji ST, Terrell KC, Huang M, et al. The Na/K-ATPase α1/Src interaction regulates metabolic reserve and Western diet intolerance. Acta Physiol (Oxf) 2021;232:e13652. 10.1111/apha.13652.

[74] Banerjee KK, Ayyub C, Sengupta S, Kolthur-Seetharam U. Fat body dSir2 regulates muscle mitochondrial physiology and energy homeostasis nonautonomously and mimics the autonomous functions of dSir2 in muscles. Mol Cell Biol 2013;33:252–64. 10.1128/MCB.00976-12.

[75] Lam TKT, Carpentier A, Lewis GF, van de Werve G, Fantus IG, Giacca A. Mechanisms of the free fatty acid-induced increase in hepatic glucose production. Am J Physiol Endocrinol Metab 2003;284:E863–73. 10.1152/ajpendo.00033.2003.

[76] Mackert O, Wirth EK, Sun R, Winkler J, Liu A, Renko K, et al. Impact of metabolic stress induced by diets, aging and fasting on tissue oxygen consumption. Mol Metab 2022;64:101563. 10.1016/j.molmet.2022.101563.

[77] Tavakkolizadeh A, Berger UV, Shen KR, Levitsky LL, Zinner MJ, Hediger MA, et al. Diurnal rhythmicity in intestinal SGLT-1 function, V(max), and mRNA expression topography. Am J Physiol Gastrointest Liver Physiol 2001;280:G209–15. 10.1152/ajpgi.2001.280.2.G209.

[78] Tavakkolizadeh A, Ramsanahie A, Levitsky LL, Zinner MJ, Whang EE, Ashley SW, et al. Differential role of vagus nerve in maintaining diurnal gene expression rhythms in the proximal small intestine. J Surg Res 2005;129:73–8. 10.1016/j.jss.2005.05.023.

[79] Stearns AT, Balakrishnan A, Rhoads DB, Ashley SW, Tavakkolizadeh A. Diurnal expression of the rat intestinal sodium-glucose cotransporter 1 (SGLT1) is independent of local luminal factors. Surgery 2009;145:294–302. 10.1016/j.surg.2008.11.004.

[80] Lanza IR, Nair KS. Muscle mitochondrial changes with aging and exercise. Am J Clin Nutr 2009;89:467S–71S. 10.3945/ajcn.2008.26717D.

[81] Short KR, Bigelow ML, Kahl J, Singh R, Coenen-Schimke J, Raghavakaimal S, et al. Decline in skeletal muscle mitochondrial function with aging in humans. Proc Natl Acad Sci USA 2005;102:5618–23. 10.1073/pnas.0501559102.

[82] Laffin M, Fedorak R, Zalasky A, Park H, Gill A, Agrawal A, et al. A high-sugar diet rapidly enhances susceptibility to colitis via depletion of luminal short-chain fatty acids in mice. Sci Rep 2019;9:12294. 10.1038/s41598-019-48749-2.

[83] Lambertz J, Weiskirchen S, Landert S, Weiskirchen R. Fructose: A Dietary Sugar in Crosstalk with Microbiota Contributing to the Development and Progression of Non-Alcoholic Liver Disease. Front Immunol 2017;8:1159. 10.3389/fimmu.2017.01159.

[84] Ghosh S, Molcan E, DeCoffe D, Dai C, Gibson DL. Diets rich in n-6 PUFA induce intestinal microbial dysbiosis in aged mice. Br J Nutr 2013;110:515–23. 10.1017/S0007114512005326.

[85] Selmin OI, Papoutsis AJ, Hazan S, Smith C, Greenfield N, Donovan MG, et al. n-6 High Fat Diet Induces Gut Microbiome Dysbiosis and Colonic Inflammation. Int J Mol Sci 2021;22. 10.3390/ijms22136919.

[86] Chan YK, Estaki M, Gibson DL. Clinical consequences of diet-induced dysbiosis. Ann Nutr Metab 2013;63 Suppl 2:28–40. 10.1159/000354902.

