## Supplementary information and figures for "Consumption of human-relevant levels of sucrose-water rewires macronutrient uptake and utilization mechanisms in a tissue specific manner"

[$] SG, TC : Equal contribution

[#] SD, BM : Equal contribution

* Address for correspondence :

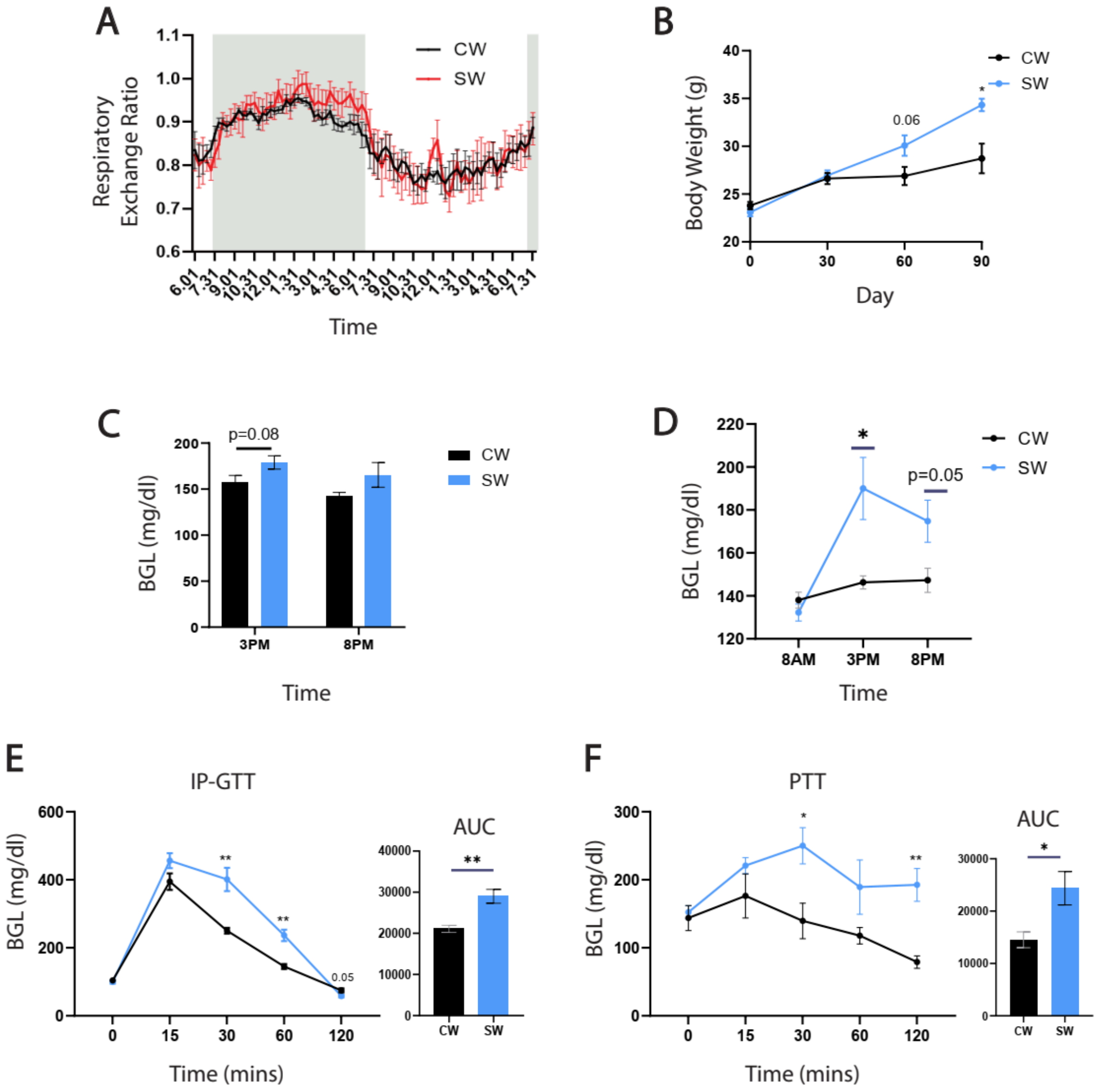


Figure S1 :

*(A) Representative respiratory exchange ratio of control and sucrose overfed C57B/6N male mice over a 24-h normal light-dark cycle. (B) Total body weights in female C57B/6N mice fed with CW and SW over 3 months (N=3, n=4-6). (C) Fasting blood glucose levels measured at 3pm, 8pm, (D) blood glucose levels measured at 8am, 3pm, 8pm measured in female CW and*

*SW mice (E) IP-GTT in overnight fasted female mice. (F) PTT in 6h fasted female mice . CW - Control mice fed with NCD + Water, SW - Sucrose overfed mice fed with NCD + 10% Sucrose soln. Experimental and technical repeats; N=3-4, n=4-6. Data represented as Mean ± SEM and analysed by t-test. P-value of 0.05 was considered significant. *P ≤ 0.05; **P ≤ 0.01; ***P ≤ 0.001.*

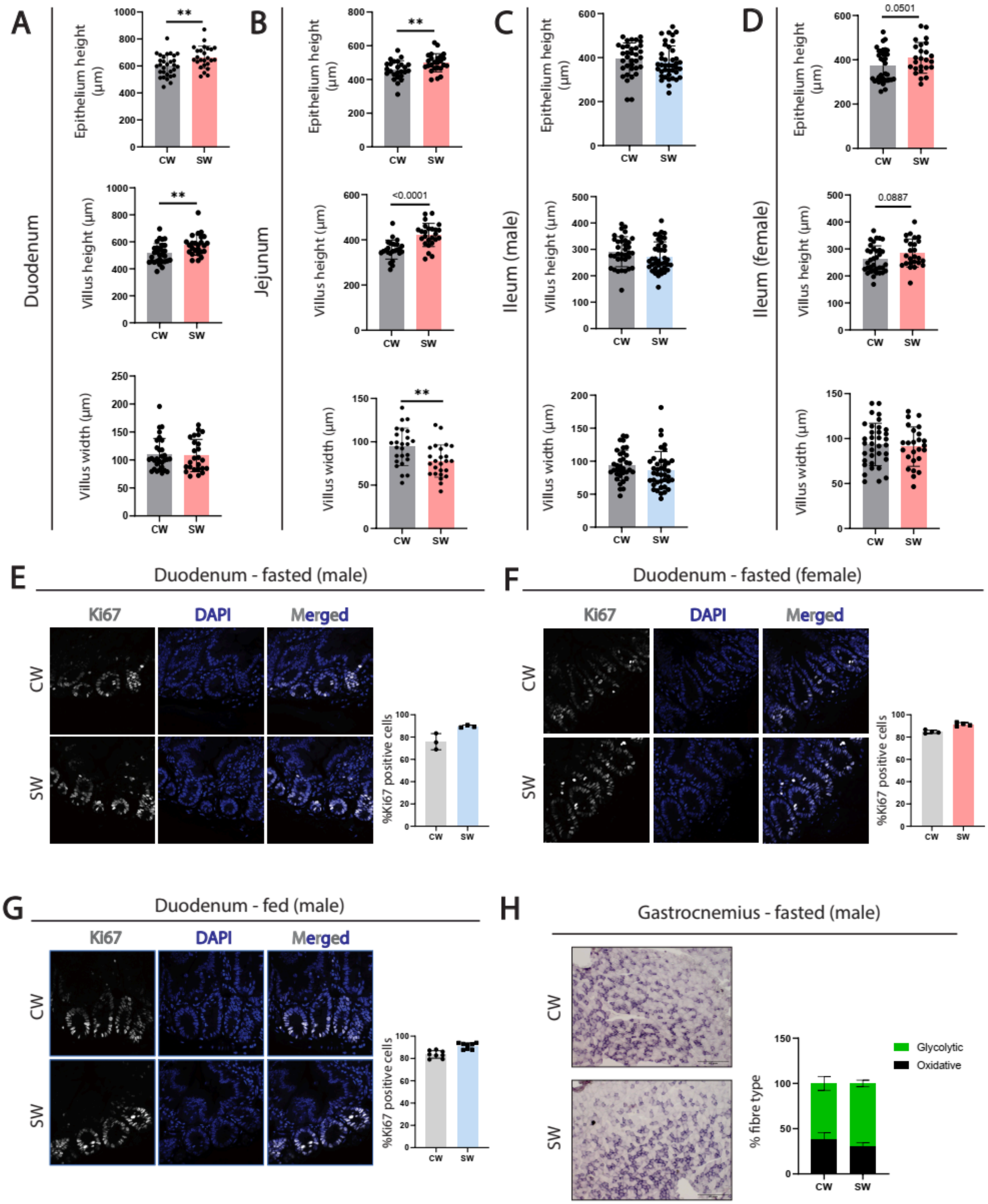
A
B
C
D
Duodenum
Jejunum
Ileum (male)
Ileum (female)
Epithelium height (µm)
Villus height (µm)
Villus width (µm)
CW
SW
**
<0.0001
0.0501
0.0887
E
Duodenum - fasted (male)
F
Duodenum - fasted (female)
G
Duodenum - fed (male)
Ki67
DAPI
Merged
%Ki67 positive cells
H
Gastrocnemius - fasted (male)
% fibre type
Glycolytic
Oxidative

Figure S2 :

*(A-D) Histological remodeling of intestine observed upon sucrose overfeeding in (A) duodenum of female mice, (B) jejunum of female mice, (C) ileum of male mice and (D) ileum of female mice. Estimations of villi parameters were done for overnight starved male (N=7-8, n=35-40) and female mice (N=5-7, n=25-35). (E-G) Representative images and quantification of proliferating cells of Duodenal crypts in (E) fasted males, (F) fasted females and (G) fed males, stained for Ki67 (grey) and counterstained using DAPI (Blue) (H) Representative sections and quantification of fibre types in Gastrocnemius muscle of CW and SW mice following overnight fasting. CW - Control mice fed with NCD + Water, SW - Sucrose overfed mice fed with NCD + 10% Sucrose soln. Data represented as Mean ± SD, and analysed by t-test. P-value of 0.05 was considered significant. *P ≤ 0.05; **P ≤ 0.01; ***P ≤ 0.001.*

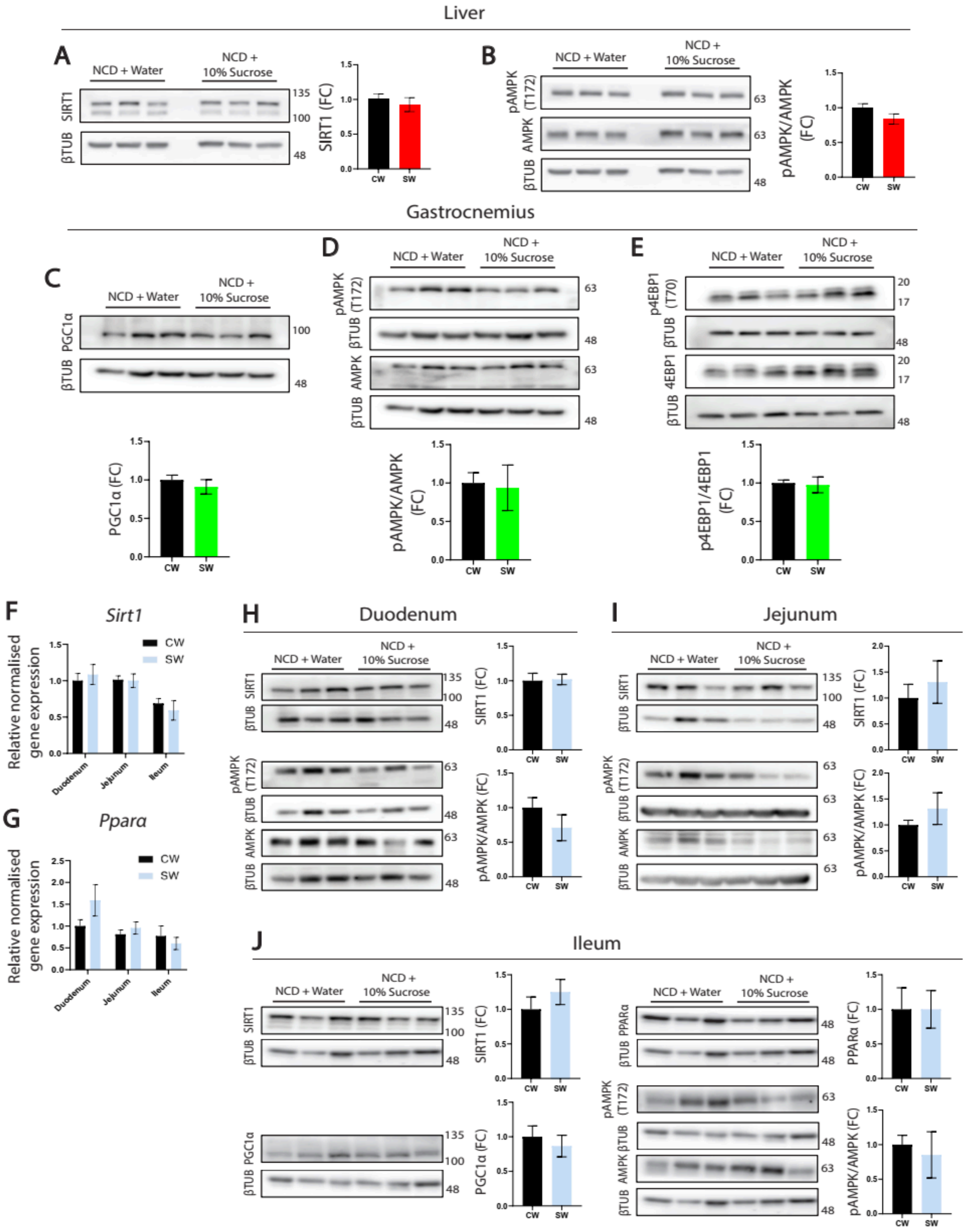

Liver
A
NCD + Water
NCD + 10% Sucrose
SIRT1
βTUB
135
100
48
SIRT1 (FC)
CW
SW
B
pAMPK (T172)
AMPK
63
pAMPK/AMPK (FC)
Gastrocnemius
C
PGC1α
PGC1α (FC)
D
E
p4EBP1 (T70)
4EBP1
20
17
p4EBP1/4EBP1 (FC)
F
Sirt1
Relative normalised gene expression
Duodenum
Jejunum
Ileum
G
Ppara
H
Duodenum
I
Jejunum
J
Ileum
PPARα
PPARα (FC)


*Figure S3 :*

*(A,B) Representative immunoblots for metabolic sensors in the lysates of liver from CW and SW mice under 12hr/fasted condition. (A) Sirt1 and (B) pT172-AMPK. (C-E) Representative immunoblots for (C) Pgc1a, (D) pT172-AMPK and (E) pT70-4EBP1 in the lysates of gastrocnemius muscle from CW and SW mice under 12hr/overnight fasted condition. (F,G) Relative intestinal (Duodenum, Jejunum and Ileum) gene expression analysis of CW and SW mice under 12hr/overnight fasted condition. (F) Sirt1 and (G) PPARa. Data represented as fold change with respect to expression in Duodenum of CW mice. (H-J) Representative immunoblots for metabolic sensors and nodal regulators of metabolism in the lysates of Duodenum, Jejunum and Ileum from CW and SW mice under 12hr/overnight fasted condition. (H) Sirt1 and pT172-AMPK in Duodenum, (I) Sirt1 and pT172-AMPK in Jejunum and (J) Sirt1, PGC1a, PPARa and pT172-AMPK in Ileum. Experimental and technical repeats; N= 2-3, n= 3. CW - Control mice fed with NCD + Water, SW - Sucrose overfed mice fed with NCD + 10% Sucrose soln. Data represented as Mean ± SEM and analysed by t-test. P-value of 0.05 was considered significant. *P ≤ 0.05; **P ≤ 0.01; ***P ≤ 0.001.*

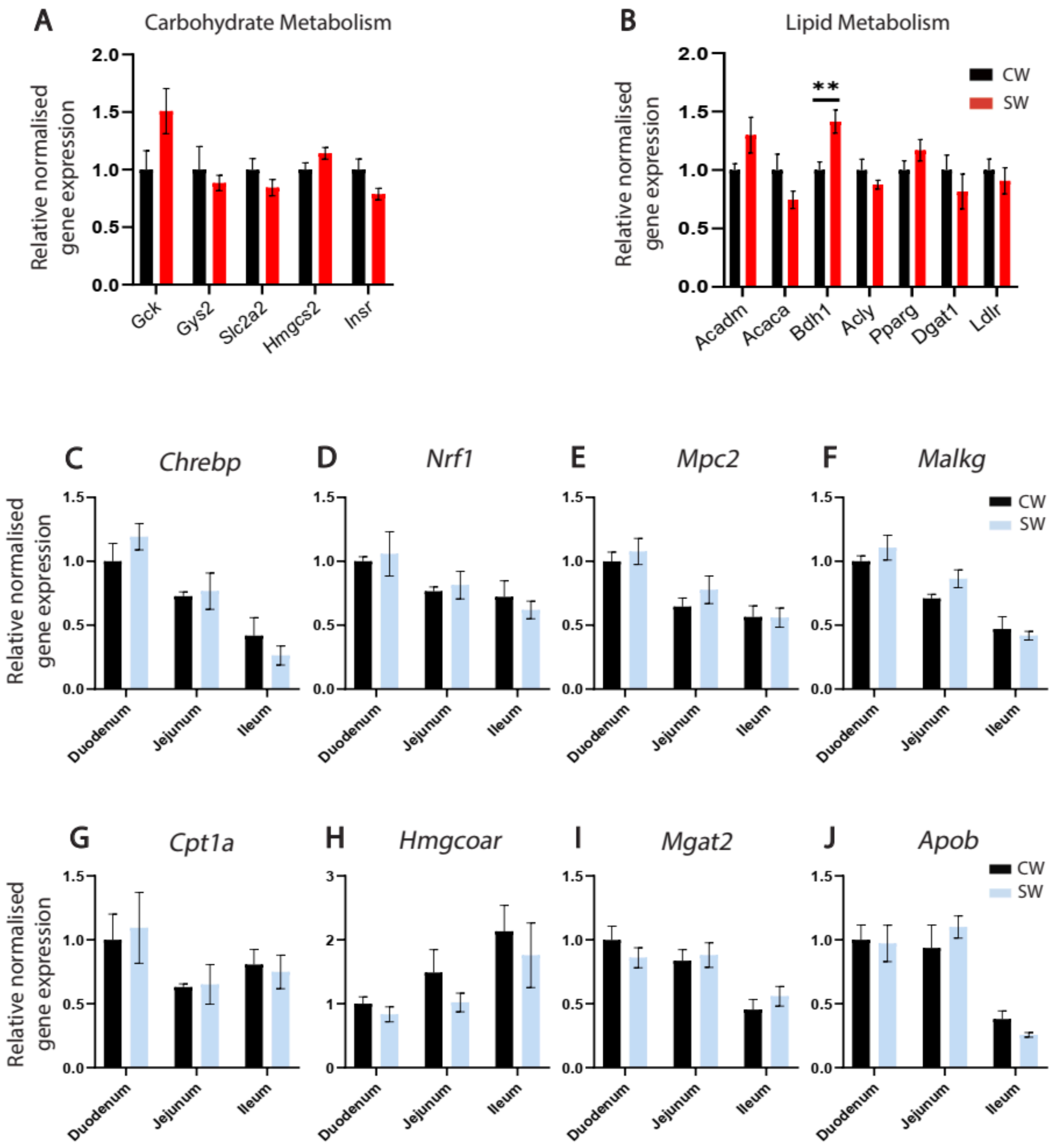
A
Carbohydrate Metabolism
Relative normalised gene expression
Gck
Gys2
Slc2a2
Hmgcs2
Insr
B
Lipid Metabolism
CW
SW
Acadm
Acaca
Bdh1
Acly
Pparg
Dgat1
Ldlr
C
Chrebp
D
Nrf1
E
Mpc2
F
Malkg
G
Cpt1a
H
Hmgcoar
I
Mgat2
J
Apob
Duodenum
Jejunum
Ileum

*Figure S4 :*

*(A,B) Relative hepatic gene expression analysis of CW and SW mice under 12hr/overnight fasted condition. (A) Carbohydrate metabolism and (B) lipid metabolism. (C-J) Relative intestinal (Duodenum, Jejunum and Ileum) gene expression analysis of CW and SW mice under 12hr/overnight fasted condition. (C) Chrebp (de-novo lipogenesis and fructose metabolism), (D) Nrf1 (biosynthesis of ETC complexes), (E) Mpc2 (mitochondrial pyruvate carrier), (F) Malkg (malate- α ketoglutarate shuttle), (G) Cpt1a (β oxidation), (H) Hmgcoar (cholesterol biosynthesis), (I) Mgat2 (monoacylglycerol to diacylglycerol conversion), (J) Apob (apolipoprotein B, LD packaging). Data represented as fold change with respect to expression in Duodenum of CW mice. CW - Control mice fed with NCD + Water, SW - Sucrose overfed mice fed with NCD + 10% Sucrose soln. Experimental and technical repeats; N=2, n=3. Data represented as Mean ± SEM and analysed by t-test. P-value of 0.05 was considered significant. *P ≤ 0.05; **P ≤ 0.01; ***P ≤ 0.001.*

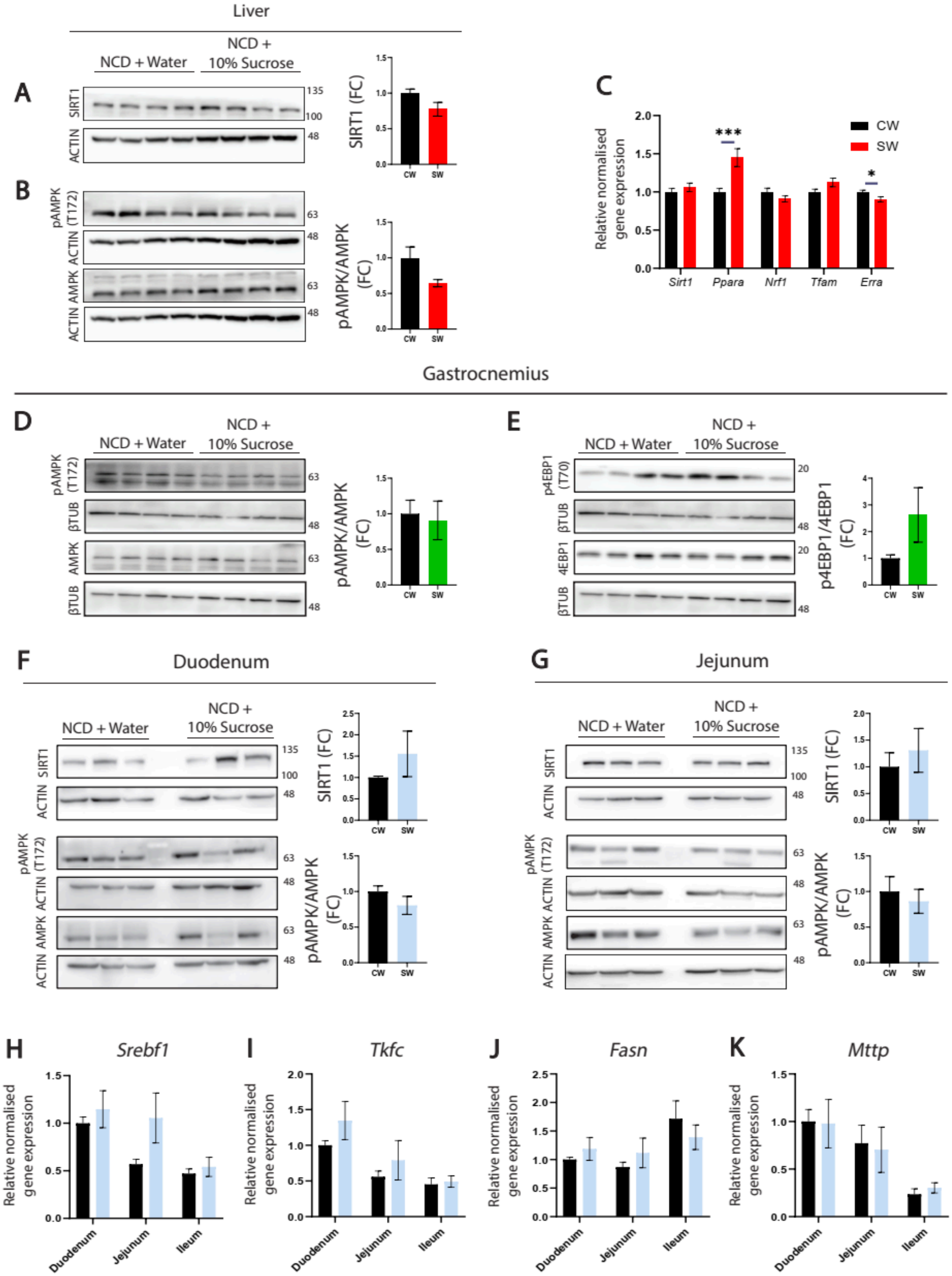

Liver
NCD + Water
NCD + 10% Sucrose
A
SIRT1
ACTIN
SIRT1 (FC)
CW
SW
B
pAMPK (T172)
ACTIN
AMPK
ACTIN
pAMPK/AMPK (FC)
C
Relative normalised gene expression
CW
SW
Sirt1
Ppara
Nrf1
Tfam
Erra
Gastrocnemius
D
pAMPK (T172)
βTUB
AMPK
βTUB
pAMPK/AMPK (FC)
E
p4EBP1 (T70)
βTUB
4EBP1
βTUB
p4EBP1/4EBP1 (FC)
F
Duodenum
SIRT1
ACTIN
pAMPK (T172)
ACTIN
AMPK
ACTIN
SIRT1 (FC)
pAMPK/AMPK (FC)
G
Jejunum
H
Srebf1
I
Tkfc
J
Fasn
K
Mttp
Duodenum
Jejunum
Ileum


*Figure S5 :*

*(A,B) Representative immunoblots for metabolic sensors (A) Sirt1 and (B) pT172-AMPK in the lysates of liver from CW and SW mice, as indicated. (C) Relative hepatic gene expression analysis for master regulators of metabolism in CW and SW mice in the fed state, as indicated. (D,E) Representative immunoblots for metabolic sensors (D) pT172-AMPK and (E) pT70-4EBP1 in the lysates of gastrocnemius from CW and SW mice, as indicated. (F,G) Representative immunoblots for metabolic sensors (F) Sirt1 and pT172-AMPK in Duodenum and (G) Sirt1 and pT172-AMPK in Jejunum from CW and SW mice, as indicated. (H-K) Relative intestinal (Duodenum, Jejunum and Ileum) gene expression analysis for (H) Srebf1, (I) Tfkc, (J) Fasn and (K) Mttp, as indicated. Data represented as fold change with respect to expression in Duodenum of CW mice. CW - Control mice fed with NCD + Water, SW - Sucrose overfed mice fed with NCD + 10% Sucrose soln. Intraperitoneal glucose was administered to the ad-libitum fed group to assess insulin responsive molecular changes in a 'postprandial state'. Experimental and technical repeats; N=2, n=3. Data represented as Mean ± SEM and analysed by t-test. P-value of 0.05 was considered significant. *P ≤ 0.05; **P ≤ 0.01; ***P ≤ 0.001.*

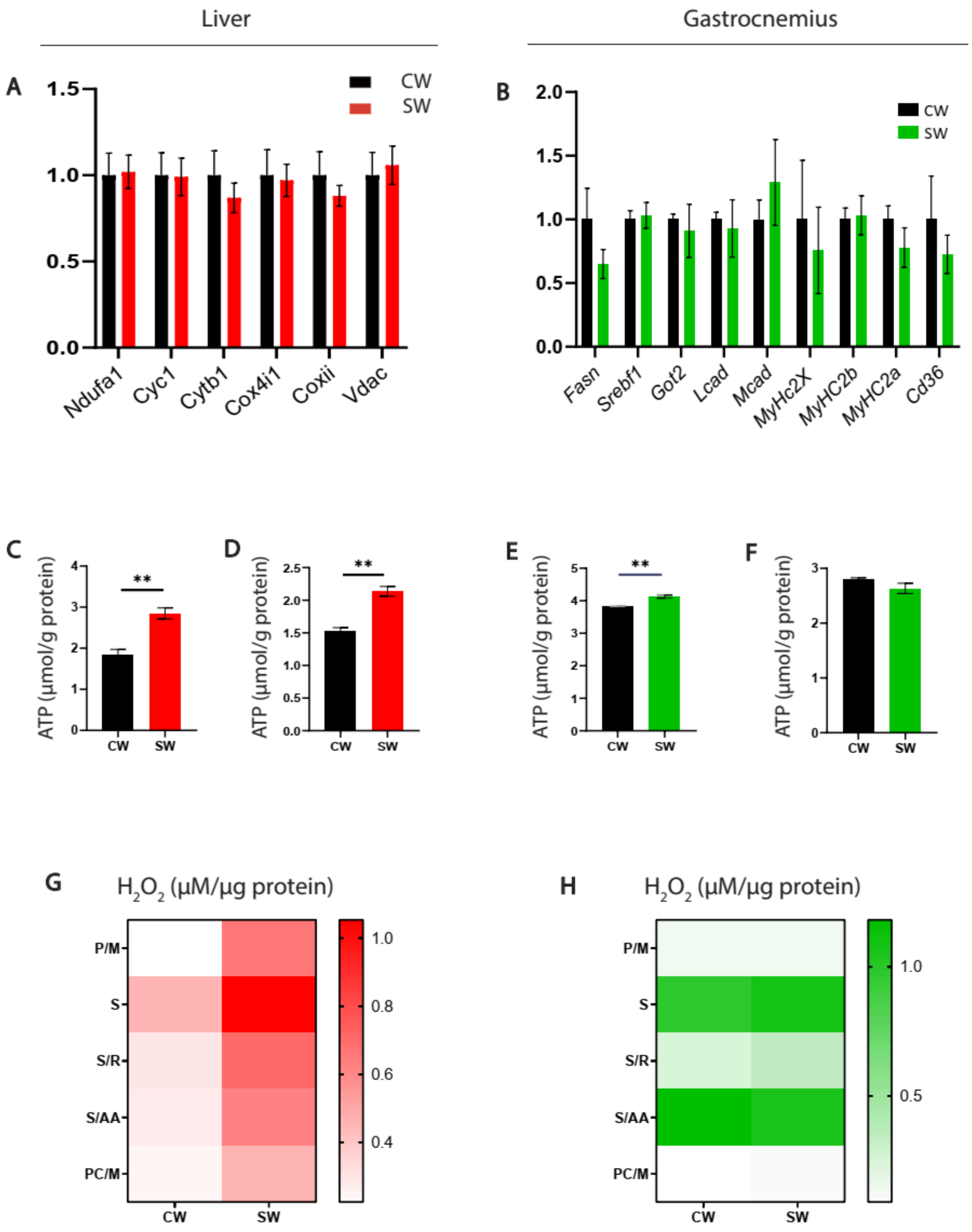
Liver
Gastrocnemius
A
CW
SW
Ndufa1
Cyc1
Cytb1
Cox4i1
Coxii
Vdac
B
CW
SW
Fasn
Srebf1
Got2
Lcad
Mcad
MyHc2X
MyHC2b
MyHC2a
Cd36
C
D
E
F
ATP (µmol/g protein)
CW
SW
**
G
H
$H_2O_2$ (µM/µg protein)
P/M
S
S/R
S/AA
PC/M

*Figure S6:*

*(A, B) Relative gene expression analysis for mitochondrial proteins assayed in (A) Liver and (B) Gastrocnemius muscle of overnight fasted CW and SW mice. (C,D) Total ATP production from mitochondria isolated from the liver in presence of (C) P/M and (D) S/R. (E,F) Total ATP production from mitochondria isolated from the quadriceps in presence of (E) P/M and (F) S/R. (G,H) ROS levels measured in mitochondria isolated from mitochondria of (G) Liver and (H) Quadriceps muscle of overnight fasted mice. CW - Control mice fed with NCD + Water, SW - Sucrose overfed mice fed with NCD + 10% Sucrose soln. Data represented as Mean ± SEM and analysed by t-test. P-value of 0.05 was considered significant. *P ≤ 0.05; **P ≤ 0.01; ***P ≤ 0.001.*

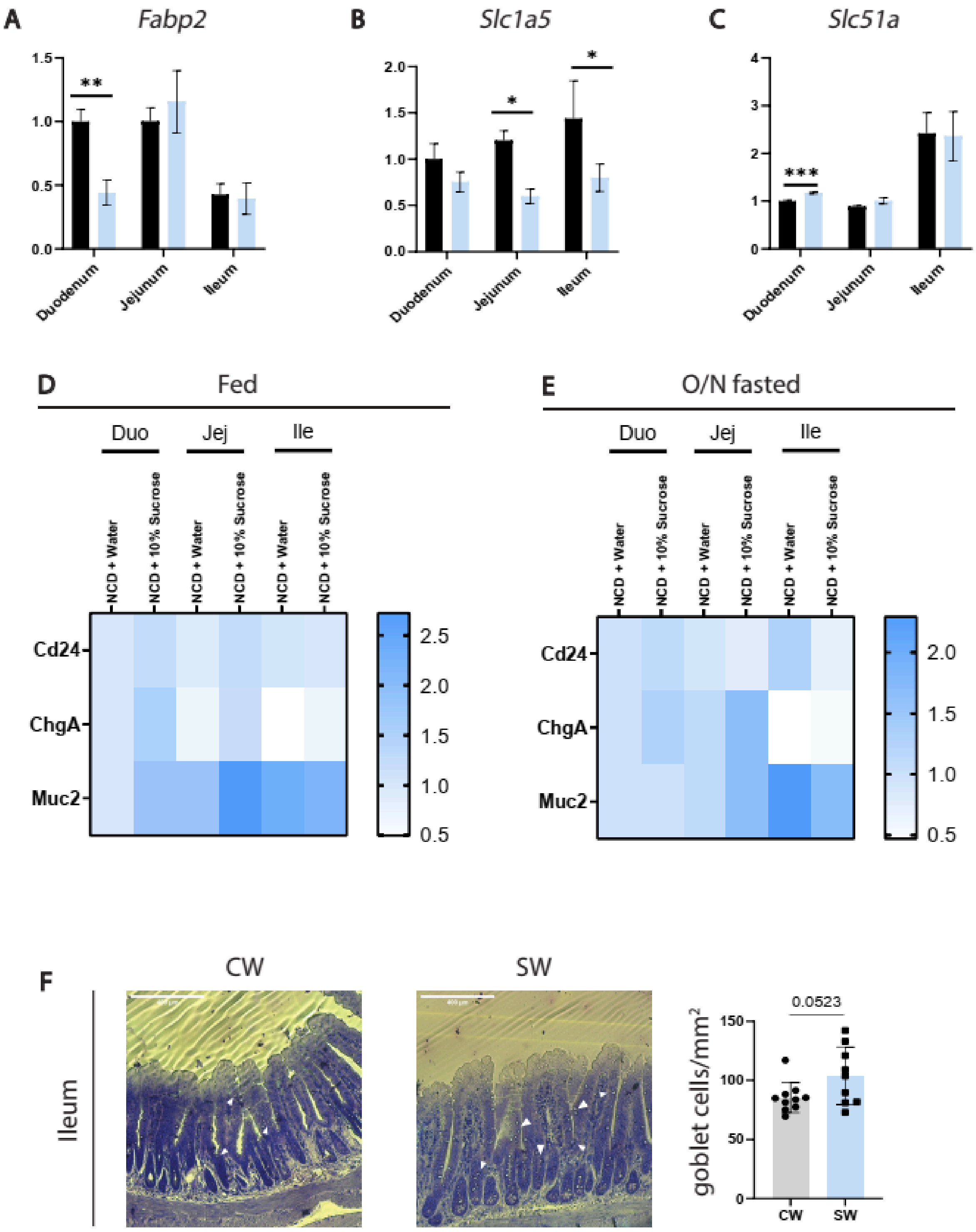
A
Fabp2
**
Duodenum
Jejunum
Ileum
B
Slc1a5
*
*
C
Slc51a
***
D
Fed
E
O/N fasted
Duo
Jej
Ile
NCD + Water
NCD + 10% Sucrose
Cd24
ChgA
Muc2
F
CW
SW
Ileum
goblet cells/mm²
0.0523

*Figure S7 :*

*(A-C) Relative intestinal (Duodenum, Jejunum and Ileum) gene expression analysis of CW and SW mice under 12hr/overnight fasted condition. (A) Fabp2 (intestinal fatty acid binding protein) (B) Slc1a5 (neutral amino acid transporter) (C) Slc51a (bile acid export pump - subunit A). (D,E) Relative intestinal (Duodenum, Jejunum and Ileum) gene expression analysis for cell-type markers in CW and SW mice under (D) fed and (E) 12hr/overnight fasted states - Cd24 (Paneth cell marker), ChgA (Enteroendocrine cell marker), Muc2 (Goblet cell marker). Data for gene expression analyses represented as fold change with respect to expression in Duodenum of CW mice. (F) Representative histological sections and quantification of goblet cell population in Ileum of sucrose overfed mice. Sections obtained from fed CW and SW mice and stained with Giemsa for visualization of goblet cells. CW - Control mice fed with NCD + Water, SW - Sucrose overfed mice fed with NCD + 10% Sucrose soln. Experimental and technical repeats; N=2, n=3-4 for gene expression analyses. Intraperitoneal glucose was administered to the ad-libitum fed group to assess insulin responsive molecular changes in a 'postprandial state'. Mean ± SEM and analysed by t-test (A-E). Experimental and technical repeats; N=2, n=8-10 for goblet cell quantification. Data represented as Mean ± SD and analysed by t-test (F). P-value of 0.05 was considered significant. *P ≤ 0.05; **P ≤ 0.01; ***P ≤ 0.001.*
